# Environment-aware genomic prediction enhances the transferability of polygenic resistance to ash dieback in *Fraxinus excelsior*

**DOI:** 10.64898/2026.04.21.719819

**Authors:** Joanna Meger, Bartosz Ulaszewski, Jarosław Burczyk

## Abstract

Ash dieback caused by *Hymenoscyphus fraxineus* threatens European ash (*Fraxinus excelsior* L.) across its range, yet natural populations retain heritable, polygenic variation in disease response. A major challenge for genomic prediction in long-lived trees is reduced transferability across heterogeneous environments, where genotype-by-environment (*G×E*) interactions may influence phenotypic expression. Here, we combined nationwide sampling across Poland (320 trees from 107 populations), whole-genome SNP data, and climate-derived predictors to test whether modelling environmental similarity and *G×E* can improve the prediction of ash dieback severity, quantified using a synthetic tree damage index (*Syn*). Environmental ordination identified a primary hydroclimatic gradient as a key driver of *Syn* (*PC1*_env_: β = 0.45 ± 0.14, p = 0.0016), although broad-scale environmental predictors explained only a modest proportion of phenotypic variance. Genome-wide association analyses revealed substantial additive genetic signal (SNP-based heritability *h*²_SNP_ = 0.63; extreme-phenotype *h*²_SNP_ = 0.81) and identified 414 suggestive loci (p < 1 × 10⁻⁵), consistent with a broadly polygenic architecture of resistance, but with pronounced local enrichment of association signals in two candidate regions on chromosomes 2 and 4. In genomic prediction, trait-enriched SNP panels consistently outperformed random panels across marker densities. Predictive ability reached *r* ≈ 0.89 in internal validation for a 500-SNP panel and remained robust (*r* ≈ 0.80) in an independent external validation set (n = 64). Incorporating *G×E* in a multi-kernel framework yielded modest but consistent gains over main-effect models, particularly under environmental extrapolation, with REML variance partitioning supported a non-zero interaction component (V*_G×E_* ≈ 14.9% and 20.9%). Our results demonstrate that ash dieback resistance is predictably polygenic and that accounting for environmental heterogeneity enhances the robustness and transferability of genomic prediction, supporting environment-aware selection and assisted migration strategies for European ash restoration.

## 1 Introduction

Tree diseases are increasingly recognized as major drivers of biodiversity loss, reduced productivity, and ecosystem disruption in forests worldwide (Boyd et al., 2013; Fisher et al., 2012; Santini et al., 2013; Seaton et al., 2025). Over recent decades, global trade, species introductions, and climate change have accelerated the emergence and spread of invasive pathogens and pests (Guégan et al., 2023; Montgomery et al., 2023). Well-known examples, such as Dutch elm disease caused by *Ophiostoma* spp. (Brasier, 1991; Gibbs, 1978), sudden oak death caused by *Phytophthora ramorum* (Rizzo & Garbelotto, 2003), chestnut blight caused by *Cryphonectria parasitica* (Anderson & Anderson, 1913; Barr, 1978), white pine blister rust caused by *Cronartium ribicola* (Geils et al., 2010; Sniezko et al., 2024), ash dieback caused by *Hymenoscyphus fraxineus* (T. Kowalski) Baral, Queloz, and Hosoya (Baral et al., 2014), and the emerald ash borer (*Agrilus planipennis*) (Poland & McCullough, 2006) illustrate how biological invasions can rapidly decimate ecologically dominant tree species, with cascading consequences for biodiversity and stability of ecosystems. Consequently, developing resilient tree populations has become a central goal in forest management and biodiversity conservation.

Genomic prediction (GP) has emerged as a promising framework for accelerating the identification of individuals with desirable traits, including disease resistance (Alemu et al., 2024). By leveraging genome-wide marker information, GP enables early estimates of breeding value without extensive phenotyping (Meuwissen et al., 2001). In major crops, GP has been widely applied to complex polygenic traits related to yield stability and pathogen resistance (Biswas et al., 2025; Guo et al., 2020; Tomar et al., 2021; Xiang et al., 2024). In long-lived forest species such as *Eucalyptus*, *Pinus taeda*, *Populus deltoides*, and increasingly *Fraxinus excelsior*, GP has shown strong potential for predicting growth, wood quality, and disease resistance (Cappa et al., 2019; Doonan et al., 2025; Guo et al., 2025; Meger et al., 2024; Resende et al., 2012c; Resende et al., 2017; Stocks et al., 2019). However, predictive accuracy often declines when models are transferred across contrasting environments or years (Bartholomé et al., 2016; Grattapaglia, 2022; Lenz et al., 2017), highlighting the need to account for environmental heterogeneity to improve predictive robustness and transferability.

Phenotypic expression of adaptive traits, including resistance to pathogens, reflects both genetic and environmental influences. Environmental heterogeneity spanning climatic gradients, soil conditions, stand structure, and local pathogen pressure can modulate trait expression and generate genotype-by-environment (*G×E*) interactions (El-Soda et al., 2014). From ecological and evolutionary perspectives, *G×E* can contribute to the maintenance of genetic diversity and play an important role in local adaptation and population persistence under ongoing environmental change (Aitken et al., 2008; Fontanari et al., 2023; Mátyás et al., 2018). In genomic prediction, ignoring *G×E* may reduce model transferability across environments and bias inference on genetic effects (Crossa et al., 2022; Millet et al., 2019; Nguyen et al., 2023; Rogers & Holland, 2022). Recent methodological advances address this challenge by modelling marker-by-environment interactions or incorporating environmental covariates through multi-environment and reaction-norm GP models (Costa-Neto et al., 2021; Crossa et al., 2022; Jarquín et al., 2014; Raffo et al., 2022). These approaches can improve predictive performance under heterogeneous conditions, particularly for traits influenced by climatic or biotic stress (Cuevas et al., 2023; Fois et al., 2021; Millet et al., 2019; Rogers & Holland, 2022). Despite this progress, *G×E*-informed GP remains underexplored in forest trees, even though long generation times, broad ecological distributions, and strong local adaptation make them especially relevant systems for studying environment-dependent genetic effects.

The European ash (*Fraxinus excelsior* L.) provides a useful system for evaluating the contribution of *G×E* to genomic prediction. Once widespread and ecologically important across Europe, *F. excelsior* has undergone dramatic declines due to the invasive ascomycete *H. fraxineus* (T. Kowalski) Baral, Queloz, and Hosoya, the causal agent of ash dieback (Baral et al., 2014; Kowalski, 2006). Since its emergence, ash dieback has spread rapidly across the continent, causing extensive mortality, altering forest structure, and affecting associated biodiversity (George et al., 2022; McKinney et al., 2014; Pautasso et al., 2013). Although susceptibility is widespread, natural populations show substantial variation in disease severity, with a small proportion of individuals displaying partial or near-complete resistance (Enderle et al., 2015; Kjær et al., 2012; Lobo et al., 2014; Muñoz et al., 2016; Pliūra et al., 2011; Stener, 2013). Recent studies indicated that resistance is heritable and polygenic, involving numerous small-effect loci (Meger et al., 2024; Muñoz et al., 2016; Stocks et al., 2019). At the same time, resistance expression varies across environmental gradients, likely reflecting differences in climate, microhabitat, pathogen pressure, and tree vigor (Havrdová, 2017; Lobo et al., 2014; Marçais et al., 2016). Together, these observations suggest that *G×E* may influence resistance phenotypes and shape evolutionary responses under continued pathogen pressure.

Although previous genomic studies have identified candidate loci and demonstrated the potential of genomic prediction for ash dieback, most models have not explicitly accounted for environmental heterogeneity, limiting transferability and obscuring environment-dependent genetic effects. Here, using a nationwide sampling design spanning over broad climatic and edaphic gradients, we integrate genomic and environmental data in a multi-kernel framework to evaluate how environmental context influences prediction accuracy in *F. excelsior*. This framework allows genetic main effects, environmental similarity, and their interaction to be modelled explicitly, providing a direct test of transferability under heterogeneous conditions. Specifically, we: (i) identify candidate SNP–trait associations for ash dieback severity using genome-wide SNP data; (ii) quantify how major environmental gradients relate to disease severity; (iii) test whether adding environmental similarity contributes to predictive performance; and (iv) evaluate whether modelling genotype-by-environment interactions via combined genomic and environmental kernels improves accuracy under environmental extrapolation and spatial transfer. Our results aim to inform environment-aware selection, breeding, and conservation strategies for European ash.

## 2 Materials and Methods

### 2.1 Plant sampling

In 2021, a nationwide survey of *F. excelsior* health status was conducted in Poland under the coordination of the Forest Gene Bank in Kostrzyca. Local foresters identified ash trees that remained free of visible ash dieback symptoms within stands heavily affected by *H. fraxineus*. These records were compiled into an internal list of candidate symptom-free trees and used to define sampling locations for the present study.

We selected 107 populations across Poland to capture broad environmental variation, including climatic gradients and site heterogeneity (Fig. 1; Table S1). In total, 320 mature trees were sampled. Where possible, three individuals per population were collected following a consistent scheme that included contrasting health phenotypes: two symptom-free trees and one severely diseased tree. Geographic coordinates were recorded for all sampled individuals.

**Figure 1.**
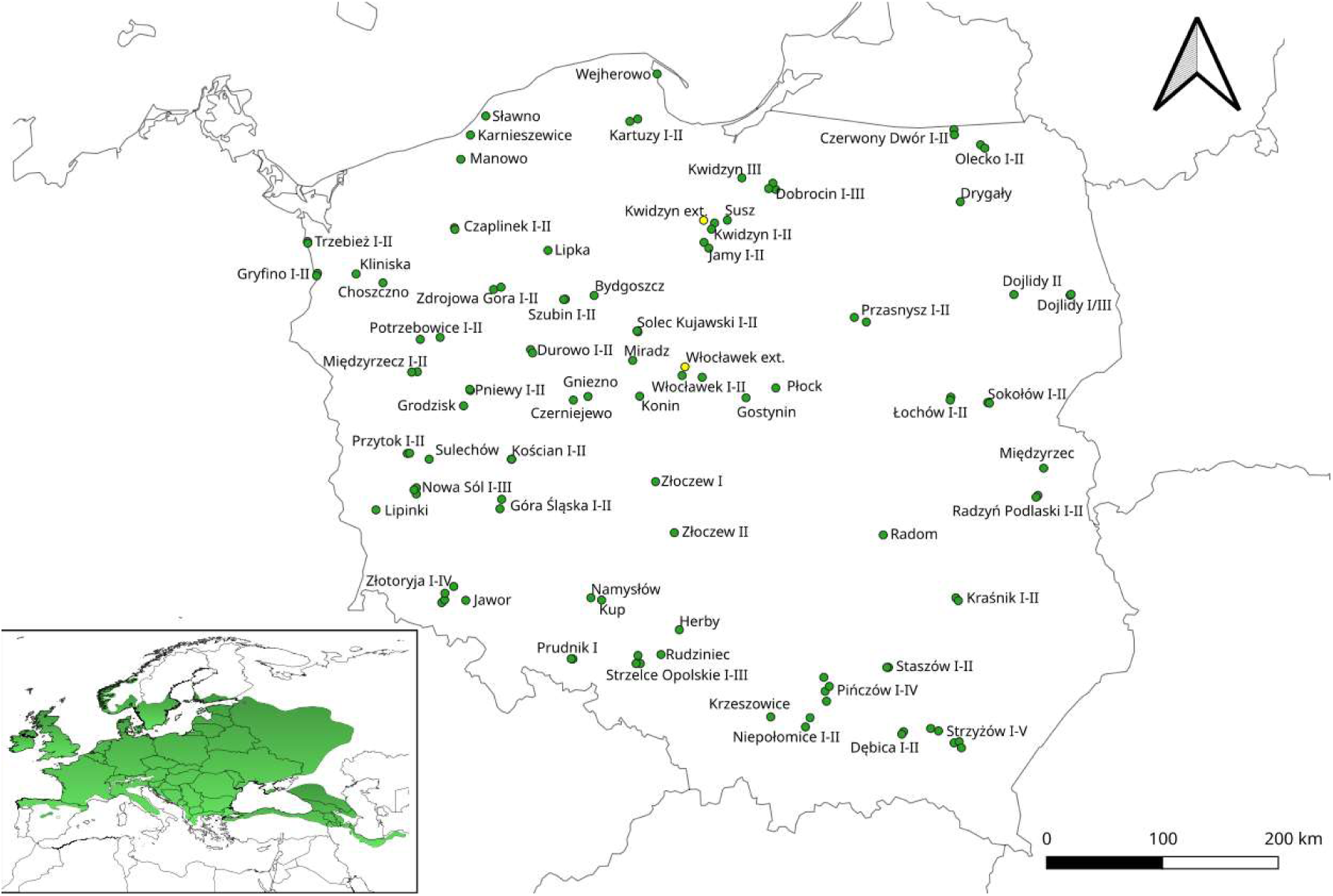
Geographic distribution of sampled *F. excelsior* populations in Poland.

Leaf material was collected in June–July 2022. Leaves were placed in paper envelopes, transported on ice, and stored at −80 °C until DNA extraction.

### 2.2 Phenotypic assessment

Tree health was quantified using the synthetic tree damage index (*Syn*), which integrates crown defoliation and crown vitality (Dmyterko & Bruchwald, 1998). For each tree, crown defoliation (Def, %) was visually estimated, and crown vitality was scored following Roloff’s classification (Roloff, 1988, 2001). Vitality scores (*Vit*) were assigned on an ordinal scale from 0 to 3, where 0 indicates an undamaged, vigorous tree; 1 a weakened tree; 2 a damaged tree; and 3 a severely damaged, declining tree. The index was calculated as:

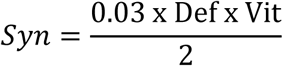

*Syn* values were treated as a continuous measure of ash dieback severity, with higher values indicating more severe crown damage and reduced vitality. Based on *Syn*, individuals were assigned to four damage classes: Class 0 (Syn ≤ 0.5; undamaged), Class 1 (0.5 < Syn ≤ 1.5; slightly damaged), Class 2 (1.5 < Syn ≤ 2.5; moderately damaged), and Class 3 (Syn > 2.5; strongly damaged). Phenotypic data for all individuals are reported in Table S1.

### 2.3 Environmental variables

To characterize local environmental conditions at each sampling location, we compiled spatially explicit climatic, drought-related, edaphic, and topographic predictors for the geographic coordinates of all sampled trees.

Climatic variables were obtained from the CHELSA v2.1 bioclimatic dataset at 30″ (∼1 km) spatial resolution (Brun et al., 2022; Karger et al., 2017). We extracted all 19 bioclimatic variables (*BIO1*–*BIO19*) describing long-term temperature and precipitation regimes. To capture additional aspects of near-surface climate, we used the CHELSA-W5E5 daily datasets (1981–2016), which provide bias-corrected fields derived from the WFDE5 and ERA5-Land reanalyses (Karger et al., 2023). From these data, we derived mean relative humidity (*hurs*), surface shortwave radiation (*rsds*), daily maximum air temperature (*tasmax*), and daily minimum air temperature (*tasmin*).

Longer-term water-balance anomalies and vegetation status were represented using CHELSA drought indices v2.1 (Chen et al., 2024, 2025). We extracted the 12-month Standardised Precipitation Index (*SPI12*) (Edwards, 1997; Mckee et al., 1993), the 12-month Standardised Precipitation–Evapotranspiration Index (*SPEI12*) (Vicente-Serrano et al., 2010), and a quality-adjusted NDVI-based greenness metric (*qkndvi*) (Chen et al., 2024, 2025).

Edaphic properties were obtained from SoilGrids 2.0 at 250 m resolution (Poggio et al., 2021) and included soil pH in water (*phh2o*), soil organic carbon (*soc*), and sand/silt/clay fractions in the 0–5 cm soil layer. Topographic variables were derived from the Shuttle Radar Topography Mission (*SRTM*) digital elevation model at 30 m resolution (Farr et al., 2007), including elevation (m a.s.l.), slope (°), and aspect (°).

Environmental predictors were first screened for association with *Syn* by calculating Pearson correlations between each predictor and *Syn* (*cor.test* in R; R Core Team, 2024). Predictors with *p* < 0.05 were retained. Within this subset, we assessed collinearity and removed redundancy by retaining one representative variable from each highly correlated group (|*r*| > 0.7) (Dormann et al., 2013). The resulting non-collinear predictor set was centred and scaled (mean 0, SD 1) and summarized using principal component analysis (PCA; *prcomp* in R; R Core Team, 2024). The first two environmental principal components were retained as synthetic covariates (*PC1*_env_ and *PC2*_env_). Ordinations were visualized using *ggplot2* (Wickham, 2016). This screening step was used to reduce dimensionality and collinearity among predictors; accordingly, the resulting environmental axes should be interpreted primarily as predictive summary variables rather than direct causal descriptors of disease environment. *PC1env* was included in downstream models as a broad-scale hydroclimatic covariate because it captured the strongest and most consistent environmental signal associated with *Syn*.

To quantify the relationship between these environmental gradients and *Syn*, we fitted a linear mixed-effects model with *Syn* as the response, *PC1*_env_ and *PC2*_env_ as fixed effects, and population as a random intercept. Models were fitted using *lme4* (Bates, 2015) and fixed-effect significance was evaluated using Satterthwaite’s approximation as implemented in *lmerTest* (Kuznetsova et al., 2017).

### 2.4 Genomic data

Total genomic DNA was extracted from leaf tissue using a modified CTAB protocol following Wang et al. (2019). DNA quantity and quality were assessed using a Quantus Fluorometer (Promega, USA) and agarose gel electrophoresis. Whole-genome sequencing (WGS) was used to obtain genome-wide SNP data for all individuals. Libraries were prepared using the TruSeq PCR-free kit (Illumina; ∼350 bp insert size) and sequenced by Macrogen Europe (Amsterdam, Netherlands) on an Illumina NovaSeq 6000 platform to generate 150 bp paired-end reads.

Raw reads were adapter- and quality-trimmed using Trimmomatic v0.33 (Bolger et al., 2014) with a 4-bp sliding window and a minimum Phred score of 30; reads shorter than 36 bp were discarded. Trimmed reads were aligned to the *F. excelsior* reference genome FRAX_001_PL (GCA_019097785.1; NCBI; Meger et al. 2024) using BWA-MEM with default parameters (Li & Durbin, 2009). Alignments were converted to BAM format, sorted, and indexed using SAMtools v0.1.19 (Li et al., 2009). Variant calling was performed in GATK v3.5 (DePristo et al., 2011). SNPs were called per sample with HaplotypeCaller (minimum Phred-scaled confidence threshold = 30) and combined into a multi-sample VCF for joint genotyping. Variants were filtered using GATK VariantFiltration and SelectVariants with thresholds QD < 2.0, MQ < 40.0, MQRankSum < −12.5, and ReadPosRankSum < −8.0, followed by additional filtering in VCFtools v0.1.16 (Danecek et al., 2011). We retained biallelic SNPs genotyped in ≥95% of individuals (site call rate ≥ 0.95), with minimum read depth ≥10 and minor allele frequency (MAF) > 0.01. For analyses requiring approximately independent markers, linkage disequilibrium pruning was performed in PLINK v2.0 using *r*² > 0.5 (Chang et al., 2015).

### 2.5 Population structure and relatedness

Population genetic structure was assessed by principal component analysis (PCA) using the quality-filtered, LD–pruned SNP dataset. PCA was performed in PLINK v2.0 (Chang et al., 2015), and eigenvectors and eigenvalues were visualized in R (R Core Team, 2024) using *ggplot2* (Wickham, 2016). The proportion of genetic variance explained by each axis was calculated from eigenvalues, and the number of informative components was evaluated using the broken-stick criterion (Jackson, 1993). To test whether sampled populations differed significantly in multivariate genotype space, we performed a permutation-based multivariate analysis of variance (PERMANOVA) using *adonis2* in vegan (Oksanen, 2020) on Euclidean distances among individuals, with 999 permutations. Pairwise genomic relatedness was estimated using a centered genomic relationship matrix (GRM) computed in GEMMA v0.98 (following the method of VanRaden, 2008) from the same filtered and LD-pruned SNP dataset (Zhou & Stephens, 2012). Kinship coefficients were visualized as a clustered heatmap using *pheatmap* in R (Kolde, 2019).

### 2.6 Genome-wide association analysis (GWAS)

Genome-wide association analyses were conducted using a univariate linear mixed model (LMM) implemented in GEMMA v0.98 (Zhou & Stephens, 2012), with the *Syn* analyzed as a continuous trait. For each SNP, the genotype at the focal marker was fitted as a fixed effect, and a random polygenic effect was modelled using the GRM (Section 2.5), thereby accounting for genome-wide additive covariance and background relatedness. Fixed-effect covariates included an intercept, the first two genetic principal components (PC1_gen_ and PC2_gen_; Section 2.5), and PC1_env_ (Section 2.3) to control for residual population structure and broad-scale environmental heterogeneity. Association significance was assessed using Wald tests, and SNP-based heritability was quantified as the proportion of phenotypic variance explained by the random polygenic effect (PVE) estimated from the fitted LMM.

Association tests were carried out on the full set of quality-filtered biallelic SNPs without LD pruning in order to retain local haplotype structure and maximize power to detect association signals. In addition to the analysis of the full dataset, including all individuals, a complementary extreme-phenotype GWAS was performed using only trees from the two most contrasting damage classes (Class 0: undamaged; Class 3: strongly damaged), applying the same LMM specification and covariates to increase phenotypic contrast. Given the moderate sample size and the expected polygenic architecture of ash dieback resistance, a suggestive genome-wide significance threshold of *p* < 1 × 10⁻⁵ was used to prioritize candidate loci for downstream analyses. This threshold was treated as a nominal and less conservative than the Bonferroni-corrected threshold based on the total number of tests (*p* < 1.19 × 10⁻⁹).

Genome-wide association results were visualized using Manhattan plots. Calibration of test statistics was evaluated with quantile–quantile (QQ) plots with 95% confidence envelopes and the genomic inflation factor (λGC), calculated from a random subset of SNPs following the genomic control framework (Devlin & Roeder, 1999). Values of λGC close to 1 were interpreted as indicating minimal residual inflation.

To place GWAS signals in a functional genomic context, SNPs exceeding the suggestive significance threshold were annotated relative to the *F. excelsior* reference genome FRAX_001_PL (GCA_019097785.1; Meger et al. 2024) and its associated gene models. Genomic coordinates were intersected with annotated features using BEDTools v2.30.0 (Quinlan & Hall, 2010) and custom R scripts, and variants were classified as exonic, intronic, untranslated regions (UTRs), or intergenic. Intergenic SNPs located within 5 kb upstream or downstream of annotated genes were additionally flagged as putative regulatory variants. Predicted functional consequences of coding variants, including synonymous, nonsynonymous, stop-gained, and start-lost mutations, were inferred using snpEff v5.4 (Cingolani et al., 2012) with a custom database built from the FRAX_001_PL genome assembly and its GFF3 annotation.

### 2.7 Multilocus genetic patterns at suggestive GWAS loci

To characterize multilocus genetic patterns underlying ash dieback severity beyond single-locus GWAS signals, we conducted post-GWAS analyses based on SNPs exceeding the suggestive significance threshold in the full-dataset GWAS (p < 1 × 10⁻⁵; 414 SNPs). To avoid overrepresentation of tightly linked variants and to focus on approximately independent genomic signals, we applied LD pruning in PLINK v2.0 (Chang et al., 2015). SNPs were pruned based on pairwise LD (r² > 0.5) within sliding windows, retaining a single representative SNP from each LD block. The resulting LD-pruned panel of 142 suggestive loci was used in subsequent multilocus analyses.

#### 2.7.1 Principal component analysis of suggestive loci

Multilocus genetic structure at the suggestive loci was examined using PCA on the LD-pruned SNP set. PCA was performed in PLINK v2.0 (Chang et al., 2015) on centred genotype dosages (coded as 0, 1, or 2 copies of the effect allele) using the *--pca var-wts* option, which returns both individual PC scores and SNP loadings. PC scores and SNP loadings were imported into R v4.4.2 (R Core Team, 2024) for downstream analyses.

Scores for PC1 and PC2 were used to visualize multilocus variation in PC space, with individuals colored by damage class (Classes 0–3). Associations between PC scores and *Syn* were evaluated using linear models of the form *Syn ∼ PC_i_* and Spearman rank correlations, implemented with the *lm* and *cor.test* functions in R.

To identify loci contributing most strongly to the primary axis of differentiation, we examined SNP loadings for PC1. Loadings were ranked by absolute magnitude, and high-contributing loci were cross-referenced with GWAS summary statistics and functional annotations. To complement the LD-pruned PCA, we also performed PCA on the full non-LD-pruned set of suggestive loci (*p* < 1 × 10⁻⁵). This analysis was used to assess the influence of local haplotypic structure on multilocus variation and to identify genomic regions in which clusters of linked SNPs contributed disproportionately to the principal axes. Results were interpreted at the regional level and used only as a complement to the LD-pruned analysis.

#### 2.7.2 Allele-frequency shifts between extreme damage classes

To test whether suggestive loci showed systematic differences in allele frequency between phenotypic extremes, we compared effect-allele frequencies between Class 0 and Class 3 individuals using an LD-pruned set of suggestive loci (Section 2.7). Genotype dosages (0, 1, 2 copies of the effect allele) were used to compute effect-allele frequency in each class (mean dosage/2). Allele-frequency shifts were quantified as *Δp = pClass3 − pClass0*; positive *Δp* values indicate that effect alleles are more frequent in strongly damaged trees. To assess concordance between allele-frequency shifts and GWAS effect estimates, we tested the relationship between *Δp* and GWAS *β* using Pearson correlation. Analyses were performed in R v4.4.2 (R Core Team, 2024) using custom scripts.

#### 2.7.3 Genotype–phenotype relationships at nonsynonymous candidate variants

To evaluate genotype–phenotype associations for coding candidates, we performed single-locus analyses for all missense SNPs detected among the suggestive loci prior to LD pruning. For each missense SNP, additive genotypes (0/1/2 effect alleles) were analyzed using linear regression with *Syn* as the response variable. Models were fitted in R v4.4.2 using *lm*, including covariates *PC1*_gen_, *PC2*_gen_, *PC1*_env_, and population to control for structure and environmental heterogeneity. Effect estimates, standard errors, and *p*-values were extracted from model summaries.

#### 2.7.4 Polygenic signal at suggestive GWAS loci

To assess whether cumulative effects of suggestive loci jointly explained variation in *Syn*, we computed polygenic scores (PGS) using all suggestive SNPs (p < 1 × 10⁻⁵) prior to LD pruning. For each individual, a raw score was calculated as the sum of effect-allele dosages weighted by GWAS regression coefficients (β) and standardized to mean 0 and SD 1. Associations between PGS and Syn were examined using Pearson correlation and linear models adjusting for *PC1*_gen_, *PC2*_gen_, *PC1*_env_, and population.

To quantify discrimination between phenotypic extremes (Class 0 vs Class 3), receiver operating characteristic (ROC) analyses were performed using PGS as the predictor, and the area under the ROC curve (AUC) was computed using pROC (Robin et al., 2011). Because SNP selection and effect-size weights were derived from the same discovery dataset, these polygenic score analyses were interpreted as descriptive summaries of within-sample multilocus signal rather than as independent measures of predictive transferability.

#### 2.7.5 External validation of multilocus structure

To assess whether the multilocus genetic signal identified in the discovery dataset was reproducible in independent material, we analyzed an external validation set comprising 64 *F. excelsior* individuals sampled in 2023 from two Polish populations (Kwidzyn and Włocławek; 32 individuals per population). None of these individuals was included in the GWAS discovery. DNA isolation, WGS, read processing, variant calling, and SNP filtering followed the same pipeline as described in Section 2.4.

Genotypes in the external dataset were restricted to the suggestive GWAS locus set identified in the discovery GWAS (414 SNPs; p < 1 × 10⁻⁵; Section 2.6). SNP identifiers and allele coding were harmonized between datasets by retaining overlapping loci and applying consistent QC and coding conventions. Genotypes were encoded as allele dosages (0, 1, or 2 copies of the effect allele).

Multilocus genetic structure was summarized by PCA computed in PLINK v2.0 (Chang et al., 2015) from the restricted SNP panel, and the proportion of variance explained by each component was calculated from PLINK eigenvalues. PCA ordinations were visualized in R v4.4.2 using *ggplot2* (Wickham, 2016) and *ggrepel* (Slowikowski, 2024). The association between PC structure and binary health status (healthy = 0, damaged = 1) was evaluated using logistic regression (*glm*, binomial family) with PC1 as the predictor. Effect sizes were reported as odds ratios (OR) with 95% confidence intervals; PC1 was also standardized (OR per 1-SD increase). Discriminative ability was quantified using ROC curves and AUC computed with *pROC* (Robin et al., 2011). An operating threshold was selected using the Youden index and summarized using sensitivity and specificity; classification outcomes were additionally summarized as counts of true positives, false negatives, true negatives, and false positives. Because PCA axes have an arbitrary sign, PC orientation was set consistently for interpretation (equivalent results are obtained after multiplying PC scores by −1).

### 2.8 Genomic prediction models

Genotypes were encoded as allele dosages (0, 1, or 2 copies of the alternative allele), consistent with an additive model (Meuwissen et al., 2001). Missing genotypes were imputed strictly within each training partition using a k-nearest-neighbours (KNN) approach implemented in the VIM package (Kowarik & Templ, 2016), following the general principle of training-only imputation. Genotype matrices were centred and scaled (mean 0, SD 1) using parameters estimated from training data only, and the same transformation was applied to validation data. All preprocessing, kernel construction, and cross-validation procedures were implemented in R v4.4.2 (R Core Team, 2024). GWAS-informed SNP rankings were recalculated within each training partition only, and the resulting top-*N* marker sets were then applied to the corresponding validation data to minimize information leakage.

To evaluate the effects of marker density and marker selection strategy on predictive performance, models were fitted using SNP panels of increasing size (100, 200, 500, 1,000, 2,000, 5,000, 10,000, 25,000, and 50,000 markers). Two complementary strategies were used to construct marker panels. First, GWAS-informed panels were generated by ranking SNPs according to association strength with *Syn* (Wald test *p*-values) and retaining the top N markers at each density level (rankings derived from the GWAS performed in this study; Section 2.6). Second, random panels of identical size were generated by sampling SNPs at random from the genome-wide LD-pruned dataset.

#### 2.8.1 Genetic main effects

To model genetic main effects, we used a hybrid prediction framework combining fixed effects for the selected SNP panel with a random polygenic background, fitted using the *sommer* package (Covarrubias-Pazaran, 2016):

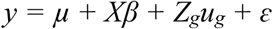

where *y* is the vector of standardized *Syn* values, μ is the intercept, *X* is the design matrix for the selected SNPs as fixed effects, *β* is the vector of marker effects, and *u_g_* is the random genomic effect, assumed to follow 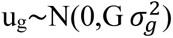), where *G* is the genomic relationship matrix computed according to (VanRaden, 2008). Residuals were assumed to be 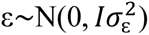).

#### 2.8.2 Environment main effects

Environmental main effects were modelled using an environmental relationship matrix (E-kernel), under the assumption that phenotypic similarity is partly driven by shared environmental conditions:

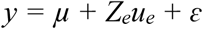

where 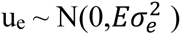). The environmental relationship matrix was computed as

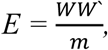

where *W* is the matrix of standardized environmental predictors, and *m* is the number of predictors included in the kernel. Environmental kernels were parameterized using BIO6, BIO15, BIO19, SPI12_08, and *PC1*_env_.

#### 2.8.3 Genotype-by-environment (*GxE*) interactions

To capture genotype-by-environment interactions, we modelled interaction covariance using the Hadamard (element-wise) product of genomic and environmental relationship matrices:

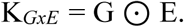

The corresponding multi-kernel mixed model was fitted using the *mmer* function in the *sommer* package (Covarrubias-Pazaran, 2016):

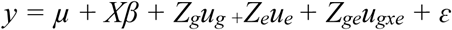

with 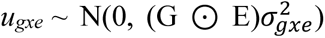). This formulation allows prediction to incorporate additive genetic effects, shared environmental effects, and environment-dependent genetic performance (Jarquín et al., 2014). Where ridge regularization of fixed SNP effects was required, it was implemented through *glmnet* (Friedman et al., 2010) or within the mixed-model framework of *sommer* (Covarrubias-Pazaran, 2016).

#### 2.8.4 Cross-validation and model evaluation

Predictive performance was evaluated under three partitioning schemes designed to assess baseline accuracy, environmental extrapolation, and spatial transfer. Model transferability was additionally assessed using an independent external validation set comprising 64 individuals (Table S9).

First, random cross-validation (CV) was used as a baseline: individuals were randomly assigned to five folds, models were trained on four folds (80%) and evaluated on the remaining fold (20%), and the procedure was repeated across 10 independent random partitions. Second, leave-one-environment-cluster-out cross-validation (LOECO) was used to test environmental extrapolation: individuals were assigned to five environmental clusters using *k*-means clustering based on *PC1*_env_ (Section 2.3), and in each fold, one cluster was withheld for validation while the remaining clusters served for training. Third, leave-one-geographic-cluster-out cross-validation (LOGCO) was used to assess spatial transfer: individuals were assigned to five spatial clusters using *k*-means clustering based on geographic coordinates, and in each fold, one cluster was withheld while the remaining clusters were used for model training.

All preprocessing and model-building steps were performed within the training data of each fold, as described above, and the resulting parameters were applied to the corresponding validation data. For internal cross-validation based on continuous *Syn* values, predictive ability was quantified as Pearson’s correlation coefficient (*r*) between observed and predicted values. Results were summarized as mean ± SD across the 10 replicates for random CV and across the five holdout folds for LOECO and LOGCO. For the external validation set, predicted *Syn* values were converted to binary classes using a threshold of 1.5 (healthy: Syn ≤ 1.5; damaged: Syn > 1.5), and agreement with observed health status was quantified using Pearson’s correlation (phi coefficient). Because the external dataset differed in phenotypic scale and structure from the discovery dataset, this analysis was interpreted primarily as a test of predictive transferability rather than as a direct replication of within-dataset performance.

#### 2.8.5 Partitioning of phenotypic variance components

To quantify the relative contributions of additive genetic, environmental, and genotype-by-environment (*G×E*) effects to variation in Syn, variance components were estimated by restricted maximum likelihood (REML) using the *mmer* function in the *sommer* package (Covarrubias-Pazaran, 2016). The same kernel structures as in the *G×E* prediction model (Section 2.8.3) were used, but the model was fitted for variance partitioning rather than for prediction.

Additive genetic variance (V*_G_*) was modelled using the genomic relationship matrix (K*_G_*), environmental variance (V*_E_*) using an environmental similarity matrix (K_E_) derived from *PC1*_env_, and the interaction variance (V*_G×E_*) using the Hadamard product (K_G_ ⊙ K_E_). Variance proportions were calculated as V_G_/V_P_, V_E_/V_P_, and V*_G×E_*/V_P_, where V_P_=V_G_+V_E_+V*_G×E_*+V_ε_. To assess robustness and transferability, the same REML framework was applied to the independent external validation set and compared with estimates obtained from the discovery dataset.

## 3 Results

### 3.1 Phenotypic variation in ash dieback severity

The synthetic tree damage index (*Syn*) spanned a wide range of values (0.075–2.93), with an average of approximately 1.1, indicating moderate overall damage among the sampled individuals. The distribution of *Syn* appeared multimodal, reflecting substantial variation in ash dieback severity across the dataset (Fig. S1). Based on the predefined severity classes, 20.3% of individuals were undamaged (Class 0, *Syn* ≤ 0.5), 45.6% were slightly damaged (Class 1, 0.5 < *Syn* ≤ 1.5), 23.4% were moderately damaged (Class 2, 1.5 < *Syn* ≤ 2.5), and 10.6% were strongly damaged (Class 3, *Syn* > 2.5).

### 3.2 Environmental gradients across sampling sites

Correlation analyses between the complete set of environmental predictors and *Syn* identified several climatic and drought-related variables significantly associated with ash health (*p* < 0.05; Table S2). Among the 19 CHELSA bioclimatic variables, precipitation of the coldest quarter (BIO19) and minimum temperature of the coldest month (BIO6) were positively correlated with *Syn*, indicating more severe crown damage at sites with wetter cold seasons and milder winter minima. By contrast, precipitation seasonality (BIO15) was negatively correlated with *Syn*, indicating an opposite relationship with disease severity. Several additional climatic variables also showed significant associations but these were excluded from downstream analyses because of strong collinearity with the main temperature— and precipitation—related gradients.

Drought-related indices—particularly SPI12 for July–September (SPI12_07, SPI12_08, SPI12_09) and SPEI12 for July–August (SPEI12_07, SPEI12_08)—were negatively correlated with *Syn*. Given that lower SPI/SPEI values indicate drier-than-normal conditions, these negative correlations suggest that stronger or more prolonged summer drought was associated with higher disease severity (higher *Syn*), whereas wetter conditions were associated with lower *Syn*. After redundancy filtering (|*r*| > 0.7), the final non-collinear subset of significant predictors comprised BIO6, BIO15, BIO19, and SPI12_08 (Table S2).

A PCA based on these standardized predictors revealed two dominant environmental gradients explaining ∼82% of total variance (PC1 = 57.8%, PC2 = 24.1%; Fig. S2). PC1 represented a composite hydroclimatic gradient driven by winter conditions and longer-term water balance, with high loadings for BIO6 and BIO19 and an opposite contribution of SPI12 (lower SPI12 indicating drier-than-normal conditions). Accordingly, higher PC1 scores were associated with milder winter minima, higher cold-season precipitation, and lower SPI12 values indicative of stronger drought, whereas lower scores were associated with colder winter minima, lower cold-season precipitation, and higher SPI12 values. PC2 captured secondary environmental variation and reflected a contrast between higher BIO6 and SPI12_08 values and higher BIO19 and BIO15 values.

To quantify the relationship between broad environmental gradients and ash dieback severity, we fitted a linear mixed-effects model with *Syn* as the response, *PC1*_env_ and *PC2*_env_ as fixed effects, and population as a random intercept. *PC1*_env_ had a significant positive effect on *Syn* (β = 0.45 ± 0.14 SE, *t* = 3.26, *p* = 0.0016), indicating that trees from sites with higher *PC1*_env_ scores tended to exhibit more severe health symptoms. In contrast, *PC2*_env_ showed no detectable effect (β = 0.10 ± 0.14 SE, *t* = 0.73, *p* = 0.48). The random population term captured a modest proportion of the variance in *Syn* (σ²_pop_ = 0.050 vs. σ_²res_ = 0.534). In a corresponding fixed-effect model without the population term, *PC1*_env_ and *PC2*_env_ together explained approximately 4% of the phenotypic variation (*R*² = 0.039), indicating that broad-scale hydroclimatic gradients contribute modestly but significantly to ash dieback severity.

### 3.3 Genotyping data

Whole-genome sequencing of the 320 sampled trees generated on average 103 million raw paired-end reads per individual (range: 52-232 million). After quality trimming and removal of low-quality bases and adapter sequences, alignment to the FRAX_001_PL reference genome resulted in an average mapping rate of 61.7% and a mean sequencing depth of 23.6× per individual (Table S3).

Initial variant calling identified approximately 165.5 million raw SNPs across all individuals. After applying GATK quality filters, 28.8 million variants remained. Subsequent filtering for bi-allelic sites, genotype call rate (≥ 95%), minimum read depth (≥ 10), and minor allele frequency (MAF) > 0.01 reduced the dataset to 26,135,993 high-quality SNPs. Linkage-disequilibrium pruning at *r*² > 0.5 further reduced this set to 11,064,930 SNPs, which were used in downstream analyses of population structure and genomic prediction.

### 3.4 Genetic variation and population structure

Genetic PCA based on genome-wide LD-pruned SNPs revealed weak but significant population structure among the 320 *F. excelsior* individuals analyzed. The first two principal components (PC1 and PC2) explained 18.1% and 13.0% of the total genetic variance, respectively, together accounting for over 31% of the observed genomic variation (Fig. S3). Most individuals formed a compact cluster, with a small number of samples displaced along PC1 and a few additional outliers along PC2, consistent with subtle geographic or demographic substructure. The broken-stick model confirmed that the leading axes captured more variance than expected under a random distribution, supporting their interpretation as informative components of population structure. A PERMANOVA (999 permutations) detected significant genetic differentiation among populations (F = 3.41, p < 0.01), consistent with subtle but non-random structure.

The genomic relationship matrix (GRM) derived from GEMMA showed very low pairwise kinship values (range = 0.006-0.097), confirming that the dataset did not contain close relatives or duplicate genotypes (Fig. S4). The distribution of relatedness coefficients was strongly right-skewed, with most values near zero, consistent with independent sampling across populations. Both the GRM and the first two genetic principal components were subsequently used to account for background relatedness and population structure in GWAS models.

### 3.5 Genome-wide association signals underlying ash dieback severity

Genome-wide mixed-model association analyses revealed a substantial additive genetic component underlying variation in ash dieback severity. In the analysis including all 320 phenotyped trees, the SNP-based heritability estimate was high (*h*²_SNP_ = 0.63), indicating that a large proportion of phenotypic variance in *Syn* can be attributed to genome-wide additive genetic effects captured by the SNP dataset. The complementary analysis restricted to extreme phenotypes (Class 0 vs. Class 3; covering 65 undamaged and 33 strongly damaged trees) yielded an even higher heritability estimate (*h*²_SNP_ = 0.81), consistent with an increased genetic signal when contrasting individuals at the tails of the phenotypic distribution.

Using a nominal genome-wide significance threshold of *p* < 1 × 10⁻⁵, the GWAS based on the full dataset identified 414 SNPs associated with *Syn* variation (Fig. 2; Table S4). These variants were distributed across nearly all assembled chromosomes, with the highest concentrations observed on chromosome 2 (212 SNPs; 51% of all associated loci), followed by chromosome 4 (31 SNPs; 7%) and chromosome 16 (24 SNPs; 6%).

**Figure 2.**
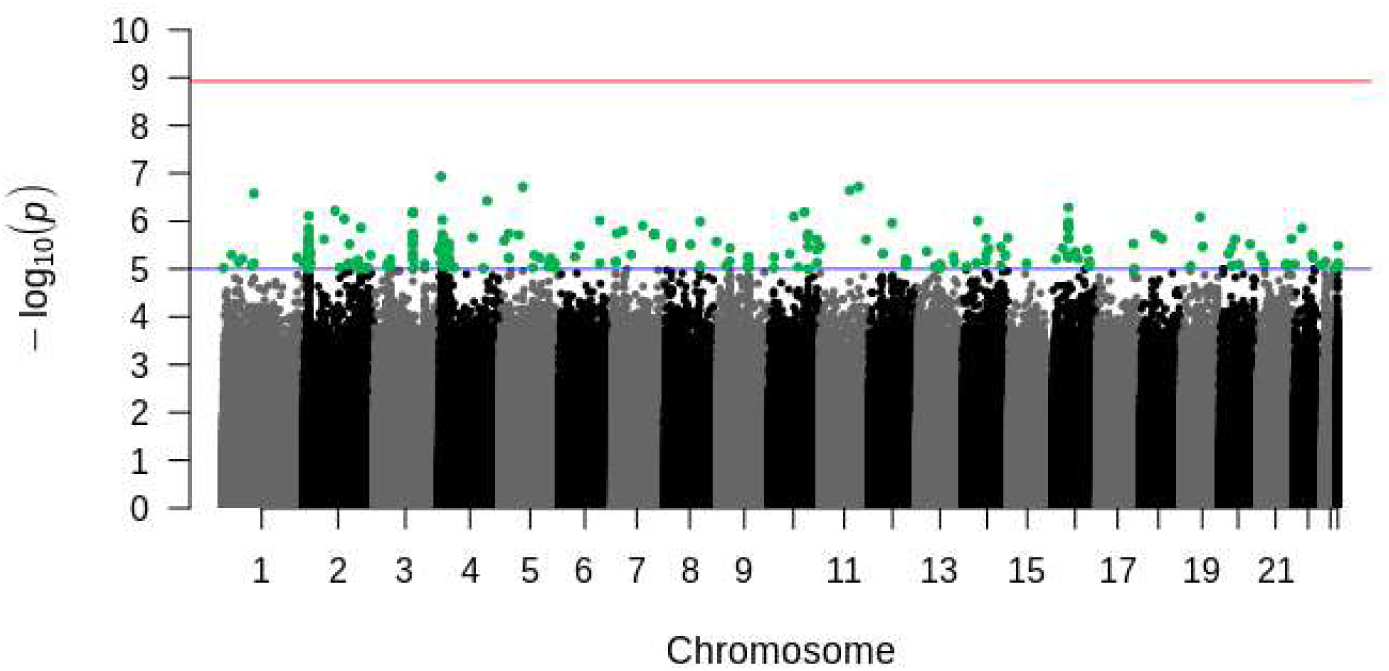
Genome-wide association results for ash dieback severity (Syn) in the full dataset. Blue and red horizontal lines indicate the suggestive (p = 1 × 10⁻⁵) and Bonferroni-corrected (p = 1.19 × 10⁻⁹) significance thresholds, respectively.

Calibration of GWAS test statistics in the full-dataset analysis was supported by QQ plots with 95% confidence envelopes (Fig. S5), which showed close agreement between observed and expected *p*-values under the null hypothesis, with minor deviation in the extreme tail. The genomic inflation factor was close to unity (λGC ≈ 1.02), indicating minimal residual inflation due to structure or cryptic relatedness.

The extreme-phenotype GWAS identified 419 SNPs exceeding the same suggestive threshold (Fig. 3; Table S5). Association signals were broadly distributed but enriched on chromosome 2 (238 SNPs; 57%) and chromosome 4 (50 SNPs; 12%). A total of 189 SNPs were shared between the two GWAS runs, supporting the reproducibility of the detected association peaks. QQ plots for the extreme-phenotype GWAS indicated appropriate overall control of test statistics (Fig. S6), with modest tail deviation (λGC = 1.11), consistent with enrichment of true signals under a polygenic architecture.

**Figure 3.**
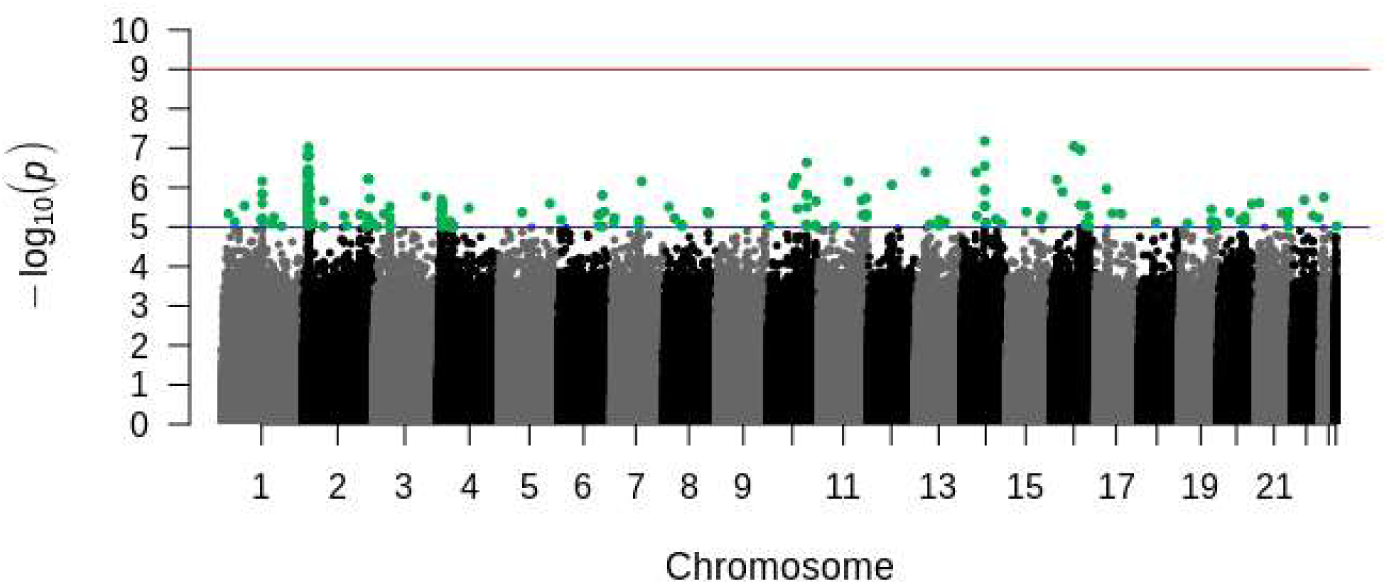
Genome-wide association results from the extreme-phenotype GWAS (Class 0 vs. Class 3). Blue and red horizontal lines indicate the suggestive (*p* = 1 × 10⁻⁵) and Bonferroni-corrected (*p* = 1.19 × 10⁻⁹) significance thresholds, respectively.

Functional annotation of suggestive loci indicated that most association signals were located in non-coding regions (Table S4). In the full-dataset GWAS, 63 SNPs mapped within annotated genes (50 intronic and 13 exonic), including 7 synonymous and 6 nonsynonymous variants, corresponding to 26 unique genes. An additional 176 SNPs were located within ±5 kb of annotated genes, while 175 were intergenic. In the extreme-phenotype GWAS, 61 SNPs fell within genes (45 intronic and 16 exonic), comprising 5 synonymous and 11 nonsynonymous variants, mapping to 16 unique genes; 197 SNPs were located within ±5 kb of genes, and 161 were intergenic.

### 3.6. Multilocus genetic structure at suggestive GWAS loci

PCA of the LD-pruned set of suggestive GWAS loci revealed a pronounced multilocus gradient aligned with ash dieback severity (Fig. 4). The first principal component (PC1) explained 28.6% of multilocus variance, while PC2 accounted for an additional 10.6%.

**Figure 4.**
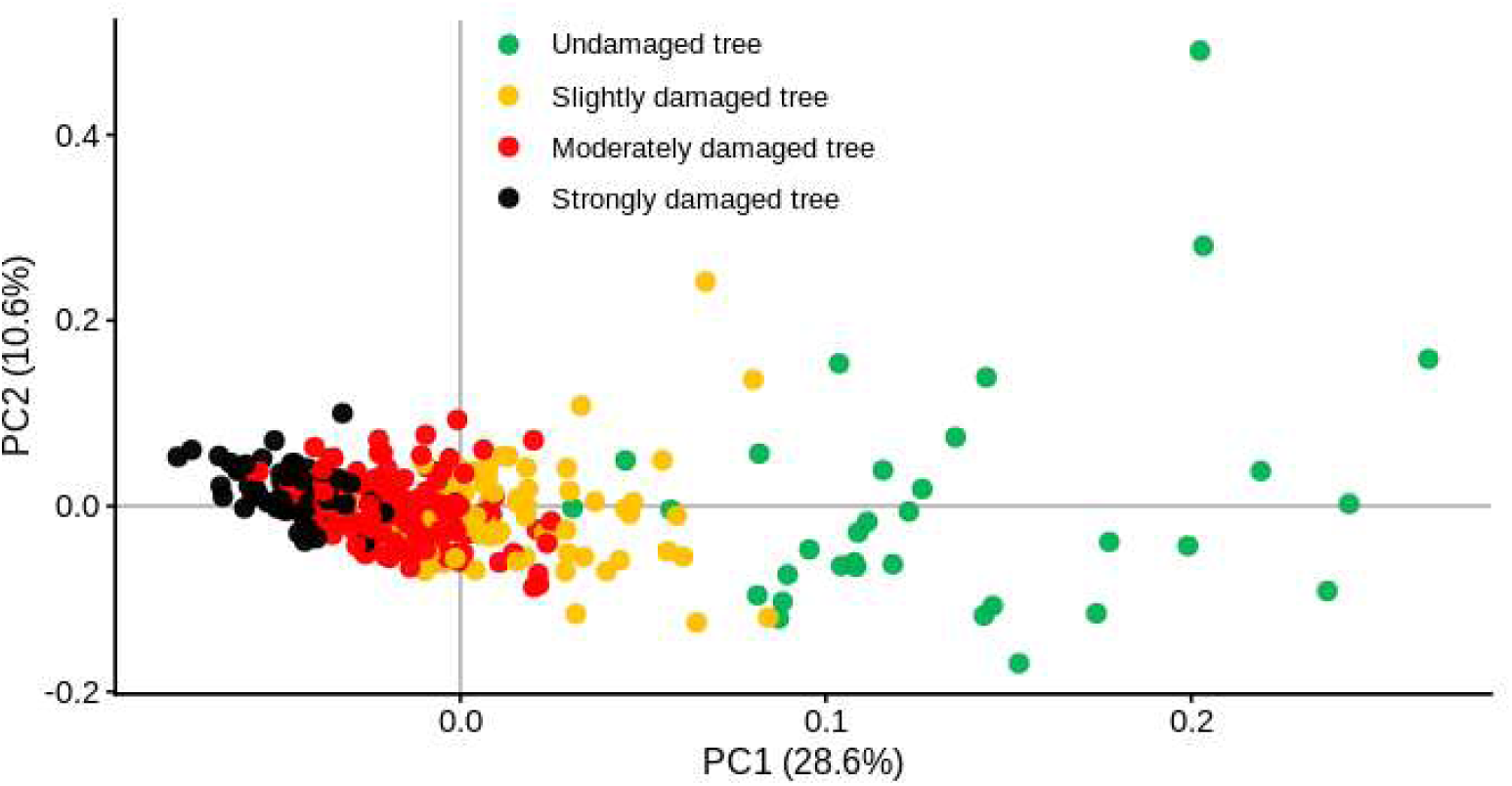
Principal component analysis of LD-pruned suggestive GWAS loci. Points represent individuals colored by ash dieback damage class.

Individuals stratified along PC1 according to disease severity, with undamaged trees (Class 0) clustering at high PC1 values and strongly damaged trees (Class 3) clustering at low PC1 values; intermediate classes occupied intermediate positions (Fig. 4). PC1 showed a strong association with *Syn* (β = −11.76 ± 0.41 SE, t = −28.6, *p* < 2 × 10⁻¹⁶), explaining 72% of phenotypic variation (*R*² = 0.72), and this pattern was supported by a strong Spearman correlation (ρ = −0.87, *p* < 2 × 10⁻¹⁶). PC2 showed only a weak and marginal association with Syn (β = 1.44 ± 0.77 SE, t = 1.87, p = 0.063) and explained <1% of phenotypic variance.

Inspection of PC1 loadings from the PCA based on the LD-pruned set of suggestive GWAS loci identified a subset of variants with high contributions to the multilocus gradient (Table S6). These top-loading SNPs were scattered across multiple chromosomes and were predominantly non-coding, most frequently intergenic or located within ±5 kb of annotated genes (upstream/downstream variants), with only a minority mapping to genic intronic regions. Among the closest annotated genes to high-loading loci were candidates involved in cellular regulation and stress-related processes, including *Tobamovirus multiplication protein 3*, *Suppressor of K(+) transport growth defect 1-like (SKD1-like)*, and a *peptide α-N-acetyltransferase*.

To assess the contribution of local LD structure to the observed multilocus pattern, we additionally examined PCA loadings from the full non-LD-pruned set of suggestive SNPs. In this analysis, the strongest PC1 loadings were concentrated almost exclusively within a dense cluster of linked variants on chromosome 2, corresponding to a prominent genomic hotspot spanning a 14.77 kb interval (3,732,327–3,747,094 bp). This region contained 158 suggestive SNPs in the *Syn* GWAS and 223 in the extreme-phenotype GWAS. Within this interval, 29 SNPs in the full-dataset analysis and 39 in the extreme-phenotype analysis mapped within the *peptide α-N-acetyltransferase* gene, whereas the remaining variants were located in flanking intergenic regions. Functional annotation identified 6 missense variants in the *Syn* GWAS and 12 missense variants in the extreme-phenotype GWAS, indicating that this region harbours both coding and nearby non-coding polymorphism. The concentration of nonsynonymous substitutions within such a narrow interval is consistent with a possible contribution of structural variation in the encoded protein, alongside linked regulatory effects, to disease response in this region.

By contrast, the strongest PC2 loadings were concentrated within a linked SNP cluster on chromosome 4 spanning a 15.95 kb interval (2,288,906–2,304,859 bp). This region contained 31 suggestive SNPs and encompassed two main candidate genes, *DETOXIFICATION 49-like* and *two-component response regulator ARR10-like*. Together, these results indicate that, although the LD-pruned PCA captured a broader multilocus gradient associated with disease severity, the non-LD-pruned ordination was strongly shaped by local haplotypic structure within a small number of candidate regions.

#### 3.6.1 External multilocus structure and phenotype association

In the independent external validation set (64 individuals; 42 healthy, 22 damaged), PCA based on the 414 suggestive GWAS loci (*p* < 1 × 10⁻⁵) captured structured multilocus genetic variation. The first two principal components explained 15.0% (PC1) and 12.4% (PC2) of total genetic variance, respectively. Healthy and damaged individuals did not form discrete clusters in PC space; instead, health status aligned with a continuous genetic gradient (Fig. 5). Damaged status was negatively correlated with PC1 (*r* = −0.46, *p* < 0.001) and more weakly with PC2 (*r* = −0.33, *p* = 0.009), indicating that higher PC1 values were associated with a lower probability of being classified as damaged. Inspection of SNP loadings suggested that several of the strongest contributors to PC1 were located near variants annotated to peptide α-N-acetyltransferase.

**Figure 5.**
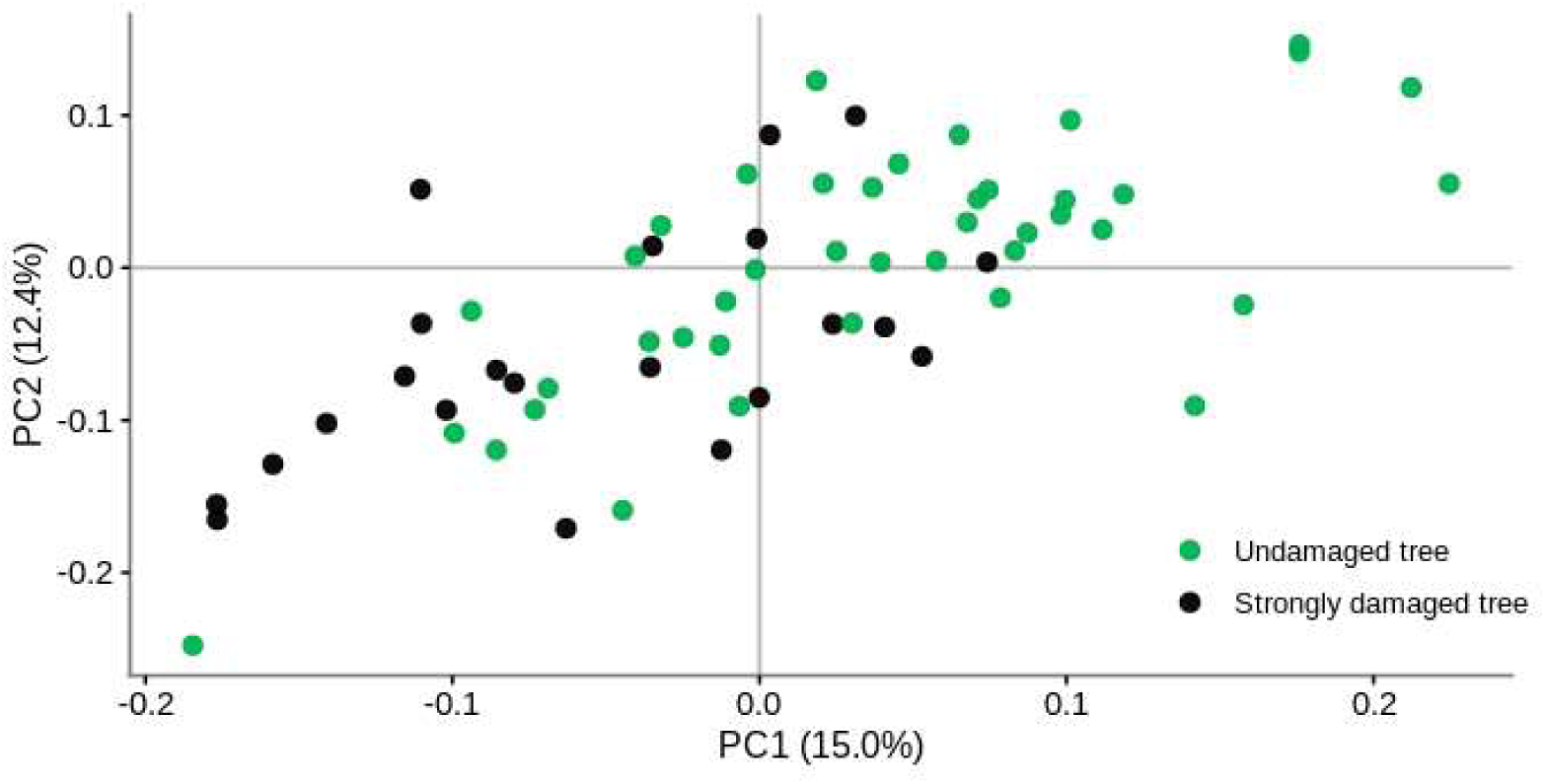
Principal component analysis of suggestive GWAS loci in the independent external validation set. Points represent individuals colored by health status.

Logistic regression confirmed that PC1 significantly predicted damaged status (p = 0.000903). When PC1 was standardized, a one–standard deviation increase in PC1 reduced the odds of being damaged (OR = 0.29; 95% CI: 0.13–0.57). Because PCA axes have arbitrary signs, ROC analysis was reported for the oriented PC1 score (equivalent to −PC1), yielding an AUC of 0.77. Using the Youden-optimized operating threshold (score = −PC1; threshold = −0.0553), classification emphasized sensitivity over specificity (0.955 vs 0.452), correctly identifying 21/22 damaged individuals (1 false negative) and 19/42 healthy individuals correctly classified (23 false positives).

### 3.7 Allele-frequency shifts between extreme damage classes

Allele-frequency differentiation between undamaged (Class 0) and strongly damaged trees (Class 3) was examined for the LD-pruned set of suggestive loci. The extreme-class comparison included 65 undamaged and 33 strongly damaged individuals. Differences in effect-allele frequency (Δ*p* = *p*Class3 − *p*Class0) were generally modest but showed a directional bias: both the median (−0.106) and mean (−0.074) were negative, indicating that the coded effect allele tended, on average, to occur at a higher frequency in undamaged trees than in strongly damaged trees. Despite this overall tendency, allele-frequency shifts were strongly concordant with GWAS effect estimates (Pearson’s *r* = 0.55, 95% CI: 0.42–0.65, *p* = 1.6 × 10⁻¹²), showing that alleles associated with increased severity in GWAS tended to be more frequent in Class 3. Detailed results are provided in Table S7.

### 3.8 Genotype–phenotype relationships at nonsynonymous candidate variants

All six nonsynonymous SNPs among suggestive loci showed significant additive associations with *Syn* after controlling for genetic structure (PC1gen, PC2gen), environmental gradients (PC1env), and population (Table S8). Effect sizes ranged from −0.87 to 0.26, with *p*-values between 1.5 × 10⁻⁵ and 2.9 × 10⁻³. For five of the six SNPs, the coded effect allele was associated with lower *Syn*, whereas one SNP showed a positive effect (β = 0.26), indicating increased damage in carriers of that allele.

### 3.9 Polygenic signal underlying ash dieback severity

Polygenic scores derived from suggestive loci were strongly associated with *Syn* (Pearson’s *r* = 0.87, 95% CI = 0.84–0.90, *p* < 2 × 10⁻¹⁶). In linear models accounting for genetic structure, environmental variation, and population effects, PGS remained a significant predictor (β = 0.41 ± 0.05 SE, *t* = 9.05, *p* < 2 × 10⁻¹⁶), and the full model explained a substantial proportion of phenotypic variation (adjusted *R*² = 0.36). ROC analysis also indicated strong within-sample discrimination between Class 0 and Class 3 individuals (AUC = 0.97). Because SNP selection and effect-size weights were derived from the same GWAS dataset, these results should be interpreted as summaries of within-sample multilocus signal rather than as independent predictive validation; transferability is evaluated separately through cross-validation and external validation in the genomic prediction analyses.

### 3.10 Genetic main effect prediction

Predictive performance depended strongly on both marker selection strategy and marker density (Fig. 6). Under internal evaluation using repeated five-fold cross-validation in the discovery population (320 individuals; 10 random partitions), models trained on GWAS-informed SNP panels consistently outperformed equally sized models trained on randomly sampled SNP panels across all tested marker densities. Predictive ability increased from *r* ≈ 0.82 at 100 SNPs to *r* ≈ 0.89 at 500 SNPs and then improved more gradually, reaching *r* ≈ 0.97 at 50,000 SNPs. By contrast, random SNP panels showed low and comparatively unstable predictive ability (*r* ≈ 0.08–0.22) regardless of marker number. In the independent external validation set, prediction accuracy increased from *r* ≈ 0.61 at 100 SNPs to a maximum of *r* ≈ 0.80 at 500 SNPs, after which it declined moderately and stabilized at higher marker densities (Fig. 6). This pattern indicates that a moderate, trait-enriched SNP panel captures most of the transferable predictive signal. In contrast, larger panels may increasingly incorporate weaker or dataset-specific signal that improves within-sample fit but generalizes less well across independent material.

**Figure 6.**
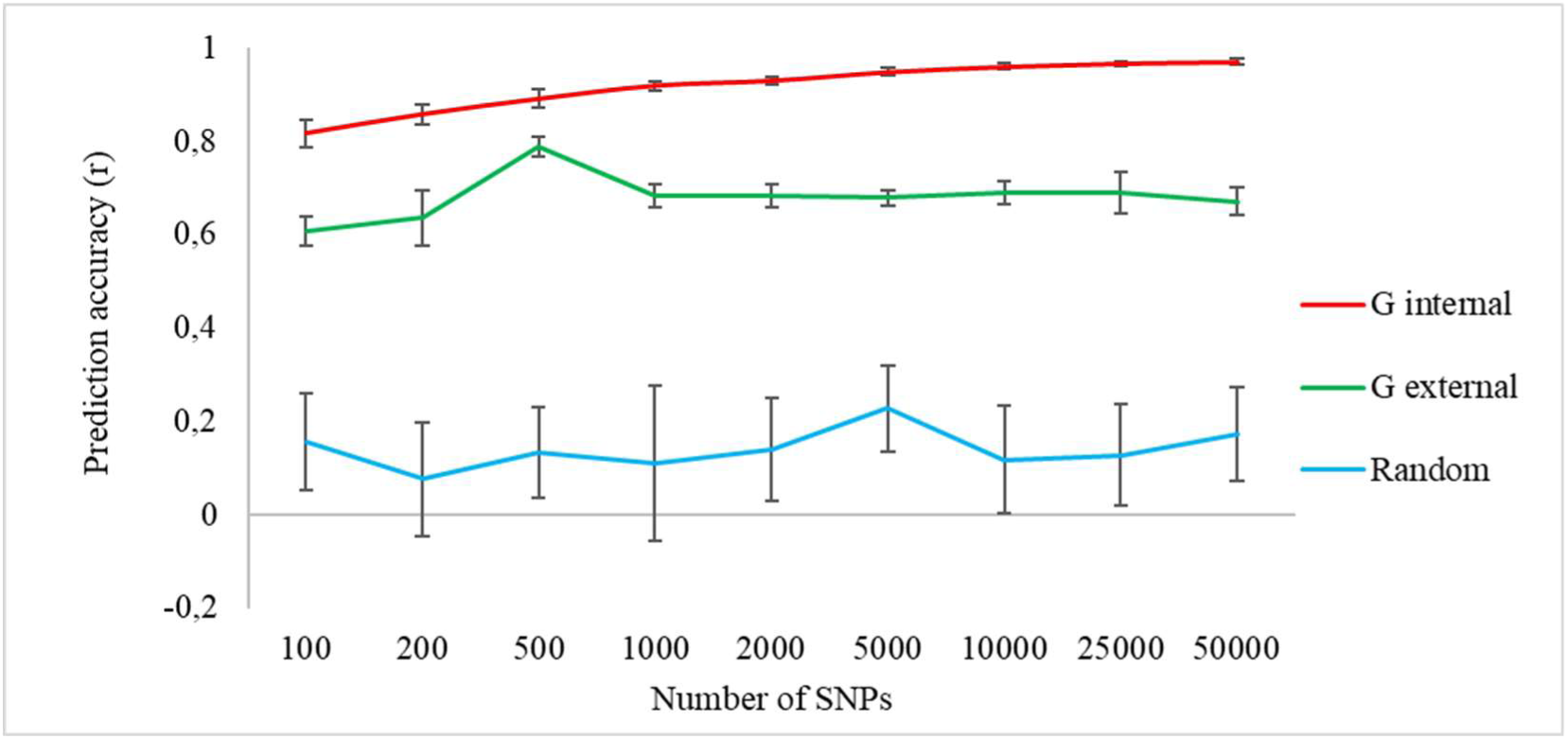
Genetic main-effect genomic prediction across marker densities and selection strategies. Predictive accuracy (*r*) as a function of SNP panel size (100–50,000). The red line (“G internal”) shows internal performance estimated by repeated five-fold cross-validation within the discovery population (n = 320; 10 replicates). The green line (“G external”) shows performance in an independent external validation set (n = 64) predicted from models trained on the full discovery dataset using the subset of SNPs shared between training and test genotype matrices. The blue line shows performance for randomly sampled SNP panels. Points represent means, and error bars indicate ±SD.

### 3.11 Environment main effect prediction

Models based exclusively on environmental similarity showed low but consistently non-zero predictive ability under random cross-validation (Fig. 7). Among single-variable models, BIO19 achieved the highest predictive ability (*r* = 0.175 ± 0.103), followed by SPI12_08 (r = 0.164 ± 0.111). BIO6 and BIO15 performed more weakly (*r* = 0.114 ± 0.132 and *r* = 0.129 ± 0.148, respectively). The strongest environmental signal was obtained for PC1_env_ (*r* = 0.192 ± 0.129), which was therefore used in subsequent *G×E* genomic prediction models.

**Figure 7.**
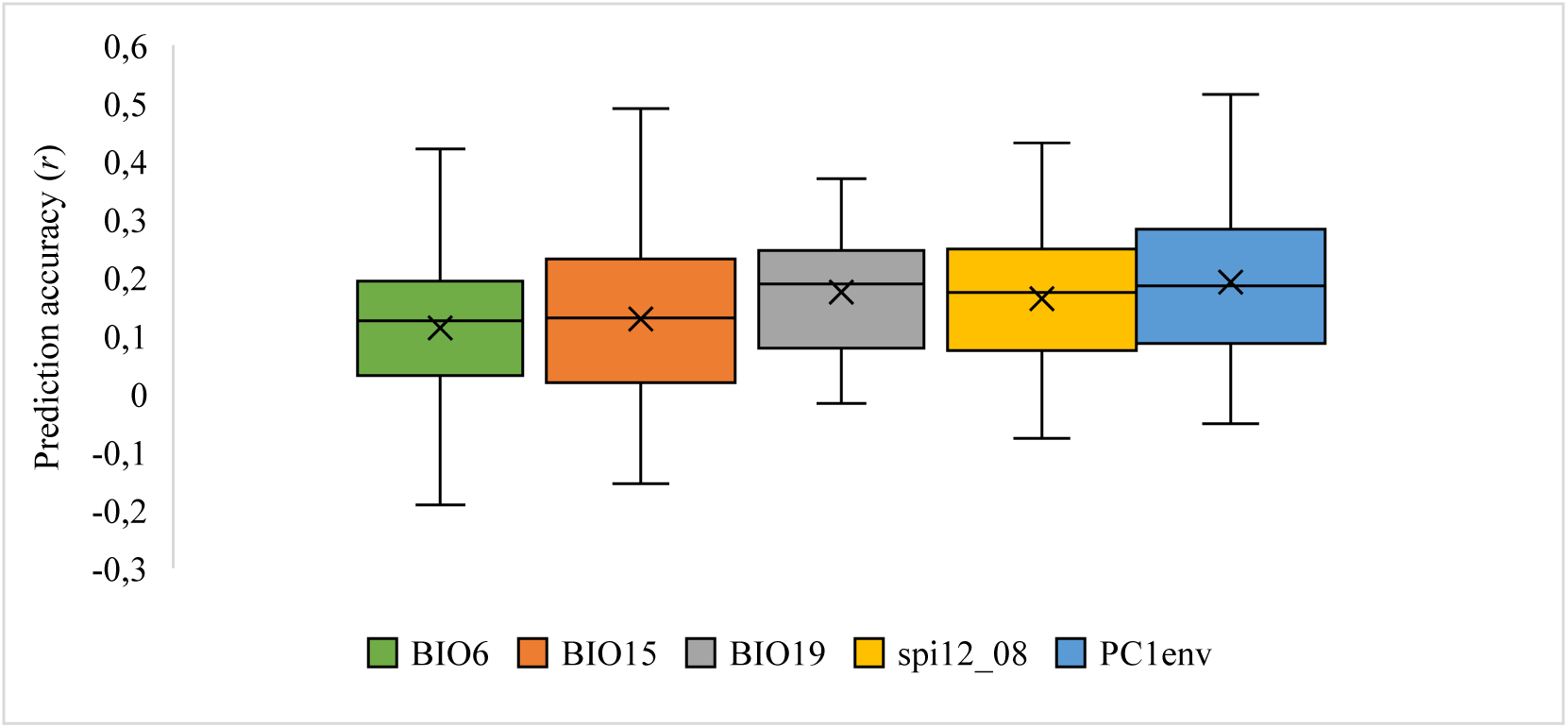
Environmental main-effect genomic prediction under random cross-validation. Distribution of predictive ability (*r*) across 10 repeated five-fold cross-validation replicates for models based solely on individual environmental predictors: BIO6 (minimum temperature of the coldest month), BIO15 (precipitation seasonality), BIO19 (precipitation of the coldest quarter), SPI12_08 (12-month standardized precipitation index), and PC1_env_ (first environmental principal component). Boxplots show the median (line) and interquartile range (box); crosses indicate means; whiskers show the spread across replicates.

### 3.12 Genotype-by-environment (*G×E*) prediction

Incorporating genotype-by-environment (*G×E*) interactions yielded small but consistent improvements in predictive ability relative to genetic main-effect models, with the magnitude of gains depending on marker density and validation scenario (Fig. 8). Under repeated five-fold cross-validation, the *G×E* model outperformed the genetic main-effect model across SNP panel sizes, with the largest differences at low marker density (Δ*r* ≈ 0.02–0.03 at 100–200 SNPs) and a smaller advantage at 500 SNPs (Δ*r* ≈ 0.015). At marker densities of 1,000 SNPs and above, performance largely converged (*r* ≈ 0.92–0.97). Variance component estimation from the multi-kernel REML model supported these patterns. In the discovery population, additive genetic effects explained most variance in *Syn* (V_G_ = 2.6648; 79.30%), the environmental main effect was small (V_E_ = 0.0196; 0.58%), and the interaction component was detectable (V*_G×E_* = 0.5007; 14.90%), with the remainder attributed to residual variance (Vε = 0.1753; 5.22%).

**Figure 8.**
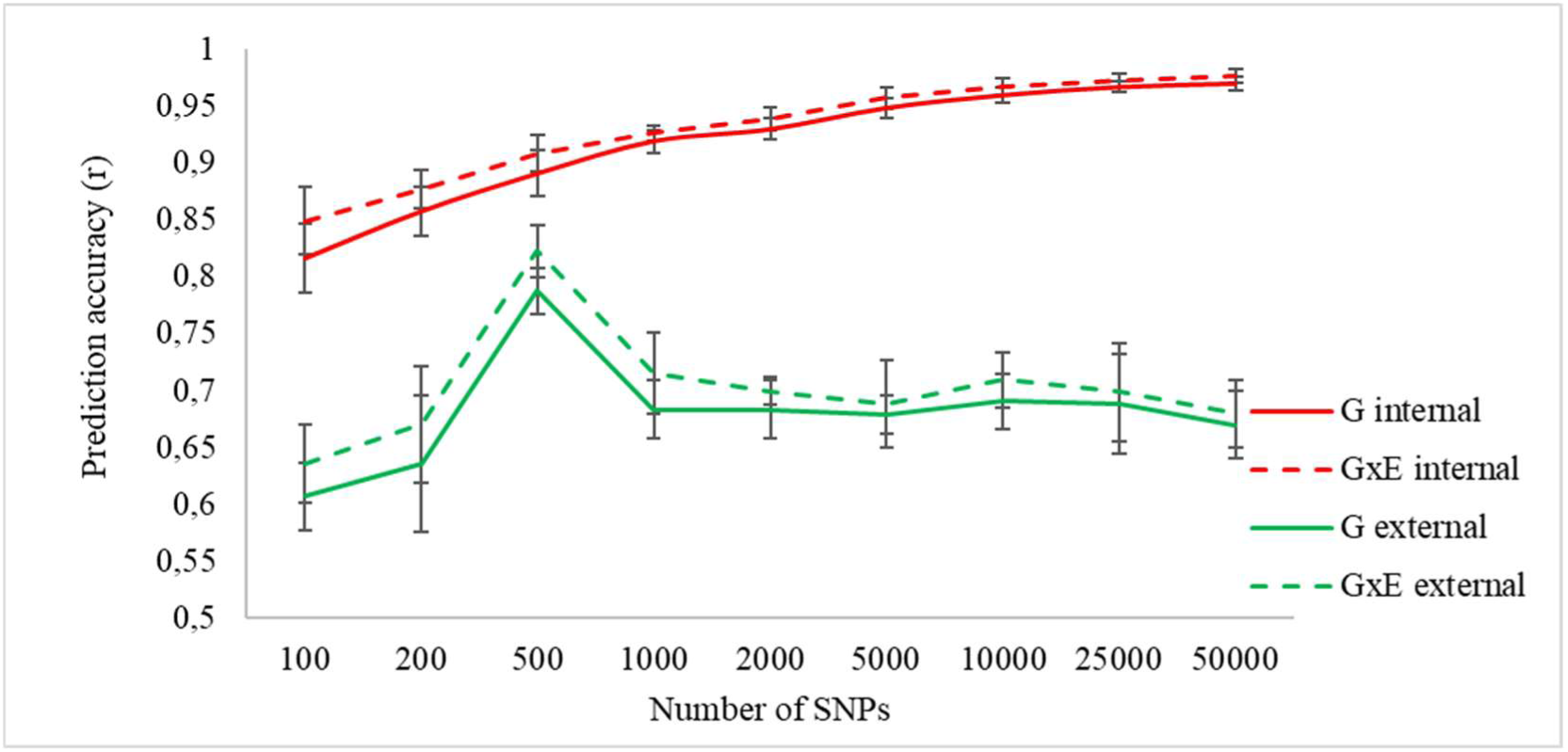
Genotype-by-environment (*G×E*) genomic prediction across marker densities. Predictive accuracy (*r*) of genetic main-effect models (G; solid lines) and models including genotype-by-environment interactions (*G×E*; dashed lines) across SNP panel sizes (100–50,000 markers). Red lines indicate internal evaluation in the discovery population (repeated five-fold cross-validation; 10 replicates). Green lines indicate independent external validation (models trained on the full discovery set and applied to external individuals using overlapping SNPs). Points represent mean predictive ability; error bars show ±SD.

In the external dataset, modelling *G×E* provided a modest but systematic improvement in transferability. Predictive ability peaked at ∼500 SNPs (r ≈ 0.80–0.82) and stabilized at r ≈ 0.68–0.71 for ≥1,000 SNPs. Variance partitioning indicated a relatively larger proportional contribution of interaction effects in the external material (V*_G×E_* = 0.0576; 20.87%), while additive genetic variance remained dominant (VG = 0.2064; 74.78%). The environmental main effect was negligible (V_E_ = 0.00064; 0.23%), and the remaining variance was attributed to residual effects (V_ε_ = 0.0113; 4.12%).

### 3.13 Model robustness and transferability across validation scenarios

Model robustness was compared across random cross-validation (CV), leave-one-environment-cluster-out (LOECO), and leave-one-geographic-cluster-out (LOGCO) schemes for the 500-SNP GWAS-informed panel (Fig. 9). Under internal validation, predictive ability declined modestly from CV (r ≈ 0.89) to LOECO (r ≈ 0.86) and LOGCO (r ≈ 0.85) for the genetic main-effect model. Incorporating *G×E* improved performance across scenarios (r ≈ 0.90 in CV, r ≈ 0.89 in LOECO, r ≈ 0.87 in LOGCO), with the largest gain under environmental extrapolation (Δr ≈ +0.03).

**Figure 9.**
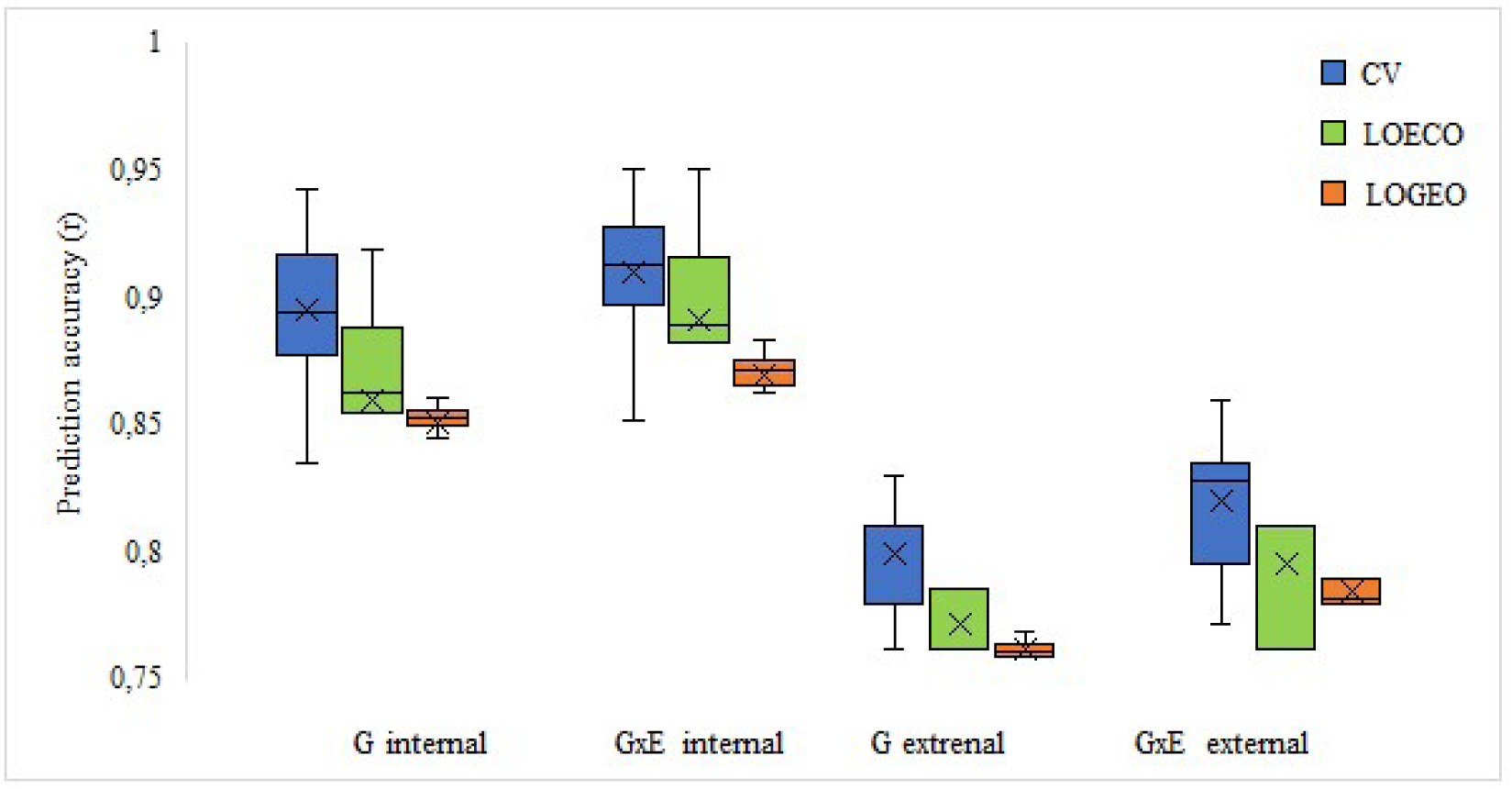
Model robustness and transferability across validation scenarios for the 500-SNP GWAS-informed panel. Predictive accuracy (*r*) of genetic main-effect (G) and genotype-by-environment (*G×E*) models for the 500-SNP GWAS-informed panel under three validation schemes: cross-validation (CV), leave-one-environment-cluster-out (LOECO), and leave-one-geographic-cluster-out (LOGCO). Internal performance was estimated via repeated five-fold cross-validation within the discovery population (10 replicates), while external performance reflects the prediction of independent individuals excluded from model training. Boxplots show the median (line) and interquartile range (box); crosses indicate means; whiskers show the spread across replicates.

In the external population, predictive ability was consistently lower than under internal validation, but the ranking remained the same (CV > LOECO > LOGCO). For the genetic main-effect model, accuracy reached r ≈ 0.80 in CV and declined to r ≈ 0.77 (LOECO) and r ≈ 0.76 (LOGCO). The *G×E* model again improved transferability (r ≈ 0.82, 0.79, and 0.78 for CV, LOECO, and LOGCO, respectively). Overall, modelling *G×E* mitigated the decline in performance under environmental and geographic extrapolation and improved robustness across validation scenarios.

## 4 Discussion

Ash dieback has created an urgent need for tools that can identify resilient *F. excelsior* genotypes and support their deployment across heterogeneous landscapes. Here, using genome-wide SNP data, broad-scale environmental covariates, and an explicit multi-kernel prediction framework, we show that variation in ash dieback severity is associated with both strong additive genetic effects and measurable environmental modulation. Across analyses, ash dieback severity exhibited a substantial SNP-based heritable component, association signals were distributed across many loci, and genomic prediction based on prioritized marker sets achieved high predictive ability. Importantly, incorporating environmental information similarity and genotype-by-environment interactions produced modest but consistent gains in predictive performance, particularly at lower marker densities and under environmental or spatial extrapolation. Together, these findings extend previous genomic work on ash dieback by showing not only that resistance is predictably polygenic, but also that its genomic prediction becomes more robust when environmental context is modelled explicitly.

### 4.1 Environmental modulation of ash dieback severity

The primary hydroclimatic gradient (*PC1*_env_) was significantly associated with ash dieback severity, although the environmental gradients considered explained only a small proportion of the observed phenotypic variation overall (4%). This is consistent with the view that disease expression in European ash is environmentally structured across multiple spatial scales: regional climate sets the broad epidemiological background, while stand structure, host density, topography, and microclimatic buffering influence how strongly that background is expressed as visible damage in individual trees (Combes et al., 2024; Cracknell et al., 2023; Enderle et al., 2019; Fuchs et al., 2024; Raykova, 2025). Accordingly, studies across Europe have repeatedly shown that ash dieback tends to be most severe in moist, sheltered environments and in stands with high ash abundance, whereas more open or drier conditions are often associated with reduced disease severity, likely owing to lower humidity, reduced inoculum build-up, and fewer opportunities for infection and lesion development (Erfmeier et al., 2019; Fuchs et al., 2024; George et al., 2022; Klesse, 2021).

Inspection of the underlying variables suggests that this signal was driven mainly by winter temperature, cold-season precipitation, and long-term summer water balance. In particular, more serious damage was associated with milder winter minima (BIO6) and wetter cold seasons (BIO19), a pattern broadly consistent with the idea that relatively moist and thermally moderate conditions may favour pathogen persistence, inoculum build-up, and infection pressure (Combes et al., 2024; Marçais et al., 2023). By contrast, lower values of SPI12_08, indicating a stronger cumulative summer moisture deficit, were also associated with greater damage, suggesting that host water stress may contribute to symptom severity in infected trees (Pliura et al., 2016). Precipitation seasonality (BIO15) contributed to the broader environmental structure, but its biological interpretation is likely less direct, as it may reflect the temporal redistribution of water availability rather than a single mechanistic driver of disease development. Taken together, these patterns suggest that ash dieback severity may be shaped by two complementary environmental pathways: (i) climatic conditions that favour pathogen development and transmission, and (ii) climatic stress that may reduce host vigour and amplify symptom expression. At the same time, the relatively small proportion of phenotypic variance explained by the environmental component indicates that broad-scale climatic variables capture only part of the environmental heterogeneity relevant to disease outcomes, much of which is likely determined by local site conditions and stand structure. This, in turn, provides a biological rationale for incorporating environmental similarity and genotype-by-environment interactions into genomic prediction models of ash dieback resistance.

### 4.2 Polygenic architecture and multilocus signals of resistance

Several lines of evidence from our study support a broadly polygenic architecture of ash dieback resistance in *F. excelsior*. First, SNP-based heritability was high in both GWAS settings (*h*²_SNP_ = 0.63 in the full dataset and 0.81 in the extreme-phenotype analysis), indicating that a substantial proportion of variation in disease severity is attributable to additive genetic effects captured by genome-wide markers. Although SNP-based heritability from mixed-model GWAS is not directly equivalent to narrow-sense heritability estimated from pedigree- or progeny-based designs, our estimates fall within the broader range reported for ash dieback-related traits. Under family, progeny, and clonal test designs, heritability of crown dieback and related symptoms is often moderate (approximately 0.37–0.53) (Kjær et al., 2012; Liziniewicz et al., 2022; Lobo et al., 2015; Muñoz et al., 2016; Pliura et al., 2016; Pliūra et al., 2011; Seidel et al., 2025), but may be substantially higher in some cases, including estimates approaching or exceeding 0.7 (Pliūra et al., 2011; Seidel et al., 2025). By contrast, heritability estimated under heterogeneous natural stand conditions may be very low, indicating that environmental noise can strongly reduce the detectable additive genetic signal (Wohlmuth et al., 2018).

Second, the genomic signal detected here was distributed broadly across the genome rather than concentrated in a single major-effect locus. In the full-dataset GWAS, 414 suggestive SNPs were identified across nearly all assembled chromosomes. with the strongest enrichment was observed on chromosome 2, followed by chromosome 4 and, more weakly, chromosome 16. The extreme-phenotype GWAS recovered a similar, though not identical, pattern, again with pronounced enrichment on chromosomes 2 and 4. Thus, resistance appears clearly polygenic but not uniformly diffused, with a genome-wide background of dispersed signal complemented by a limited number of more prominent regions of enrichment. This pattern is consistent with a polygenic architecture in which both diffuse small-effect variation and localized clusters of linked variants contribute to phenotypic differentiation. This interpretation agrees well with previous genomic studies in *F. excelsior*. Stocks et al. (2019) reported thousands of associated SNPs distributed across the genome, whereas Meger et al. (2024) found relatively few individually significant hits but strong multilocus predictability. More recent whole-genome analyses across European populations likewise showed that ash dieback crown damage is polygenic and underpinned by a widely distributed set of largely unlinked SNPs, reinforcing the view that much of the relevant signal is dispersed across the genome rather than concentrated in a single major region (Doonan et al., 2025). This interpretation is also consistent with recent pangenome analyses, in which localized signals associated with lower susceptibility were likewise interpreted as components of a broader polygenic background rather than as evidence for a single dominant resistance region (Wood et al., 2025).

In interpreting these GWAS results, it is important to acknowledge that the significance of individual associations was likely constrained by limited statistical power. Although our sampling captured broad geographic and environmental variation, the total sample size remains modest relative to the highly polygenic nature of ash dieback resistance and the vast number of markers tested. Furthermore, the *Syn* index - as a composite field-based metric – inherently carries some observer subjectivity and phenotypic noise. Together, these factors likely reduced the power to detect individual SNP effects; thus, the moderate significance of single-locus associations should be viewed as an expected feature of GWAS for complex traits, rather than evidence against a meaningful genetic basis.

Our multilocus analyses provide a more nuanced perspective. PCA of the LD-pruned suggestive loci revealed that the primary multilocus axis (PC1) was strongly correlated with *Syn*, capturing the main gradient of phenotypic differentiation. In contrast, PC2 showed only a marginal association with disease severity and explained little additional variation. While high-contributing loci identified from PC1 loadings were distributed across multiple chromosomes, analysis of the full non-LD-pruned set of suggestive SNPs showed that the strongest PC1 signal was concentrated almost exclusively within a dense hotspot on chromosome 2, spanning a ∼15 kb interval. This region exhibited a high density of associated variants, including missense mutations and nearby non-coding polymorphisms, suggesting a combination of coding and linked cis-regulatory effects. Conversely, PC2 loadings highlighted a second localized cluster on chromosome 4, encompassing *DETOXIFICATION 49-like* and *ARR10-like* genes (Xie et al., 2018; Zubo et al., 2017), though its overall phenotypic contribution was weaker. Collectively, these results indicate that ash dieback resistance is not a purely diffuse infinitesimal trait, but rather a broadly polygenic architecture where a genome-wide background of small-effect variants is punctuated by a limited number of localized regions that contribute disproportionately to disease resistance.

Across the full set of suggestive loci, genomic annotation was consistent with a predominantly non-coding and likely regulatory component of ash dieback resistance. Most associated SNPs were intronic, intergenic, or located within ±5 kb of annotated genes, whereas only a minority represented coding changes. This predominance of non-coding and near-genic signal is consistent with previous ash studies, including whole-genome and pangenome analyses, which indicate that a substantial fraction of the genomic basis of ash dieback resistance lies outside protein-coding sequence, consistent with an important role for regulatory and other non-coding variation (Doonan et al., 2025; Meger et al., 2024; Stocks et al., 2019; Wood et al., 2025). At the same time, the genes located near suggestive loci point to several biologically plausible functional categories, including defense signaling, pathogen perception, and broader stress-response pathways, although the strength of support is clearly uneven. Among previously highlighted ash candidates, *MACPF domain-containing CAD1-like* remains one of the more plausible loci, a defence-related locus previously implicated in ash dieback resistance (Stocks et al., 2019), while other genes identified here, including *probable disease resistance protein At1g61310*, *calmodulin-binding protein 60 A-like* (Kumari et al., 2022; Wang et al., 2009; Yu et al., 2025), and sphingoid long-chain bases kinase 2 (Magnin-Robert et al., 2015; Seo et al., 2021), are also plausible because related genes in other plant systems are involved in *NLR-mediated recognition effector-triggered immunity* (Chen et al., 2022), salicylic-acid signaling (Mishra et al., 2024; Peng et al., 2021), or sphingolipid-associated immunity (Saucedo-García et al., 2023; Zeng & Yao, 2022). Nevertheless, these findings should be interpreted with caution: the identified regions are best regarded as functional hypotheses rather than established mechanisms, and the observed clusters may reflect causal variants, linked regulatory polymorphisms, or local haplotypic structure.

Finally, our post-GWAS multilocus analyses provide an additional and independent line of support for a polygenic architecture of ash dieback resistance. PCA of the LD-pruned suggestive loci revealed a pronounced multilocus gradient associated with *Syn*, with healthier and more strongly damaged trees occupying opposite ends of the primary axis. Together with the directional allele-frequency shifts observed between extreme damage classes, this indicates that the GWAS signal is coherent at the multilocus level and reflects coordinated allelic differentiation related to disease severity, rather than a loose collection of isolated marker–trait associations. The external validation set further strengthened this interpretation: in independent material, health status again aligned with a continuous genetic gradient defined by the suggestive loci, and the first PCA axis significantly predicted damaged versus healthy classification. This pattern is consistent with resistance being shaped by the cumulative effects of many alleles, such that individuals vary along a continuum of multilocus susceptibility rather than separating into discrete resistant and susceptible classes. Collectively, these results indicate that the multilocus structure identified in the discovery dataset captures biologically relevant signals that are reproducible in independent material. This reinforces the conclusion that ash dieback resistance in *F. excelsior* is governed by cumulative effects distributed across many loci, while also highlighting two localized candidate regions that stand out against this broader polygenic background.

### 4.3 Genomic prediction and marker prioritization

Prediction performance was consistently higher for GWAS-informed SNP sets than for random SNP sets of the same size across all tested panel sizes, indicating that genomic prediction of ash dieback severity in *F. excelsior* is driven primarily by trait-relevant multilocus signal rather than by background genomic covariance captured by untargeted markers. Our results further show that predictive performance depends not only on which markers are used, but also on how many are retained. Under internal validation, predictive ability increased steeply at low marker densities, rising from approximately 0.82 with 100 SNPs to approximately 0.89 with 500 SNPs, and then increased more gradually towards a near-saturation plateau (0.97 at 50,000 SNPs). Independent external validation, however, was particularly informative: prediction accuracy increased from approximately 0.61 at 100 SNPs to approximately 0.80 at 500 SNPs, then declined moderately and stabilized at higher marker densities. This pattern suggests that a moderate, trait-enriched marker set may provide the best balance between capturing transferable signal and avoiding the inclusion of weaker, redundant, or dataset-specific effects that improve within-sample fit but generalize less well across independent material.

This interpretation closely aligns with previous work on European ash. showed that genomic prediction based on a relatively small set of highly informative loci could classify tree health with over 90% accuracy using 200 top-ranked SNPs, demonstrating that ash dieback susceptibility can be predicted efficiently from GWAS-enriched marker subsets. Likewise, Meger et al. (2024) found that GWAS-prioritized SNP panels achieved markedly higher accuracy than random subsets, with high predictive performance across panels of several hundred to several thousand SNPs. Recent whole-genome analyses across European ash populations reinforce the same conclusion: Doonan et al. (2025) reported that as few as 100 unlinked SNPs ranked by association strength explained 55% of the variation in ash dieback tolerance, increasing only modestly to 63% when the marker number rose to 9,155. Taken together, these studies indicate that for ash dieback, much of the predictive signal is concentrated in a relatively small subset of trait-enriched loci, and that expanding marker density beyond this core set may yield diminishing returns in terms of transferability.

### 4.4 Genotype-by-environment interactions contribute to predictive transferability

Genomic prediction in forest trees often achieves high apparent accuracy when training and test sets are closely related, yet performance commonly declines when models are applied across families or populations, and also across years, sites, or deployment zones (Beaulieu et al., 2014; Lauer et al., 2022; Lenz et al., 2017; Resende et al., 2012c). This pattern suggests that predictive ability in many datasets is partly contingent upon realized relatedness and underlying family or population structure; consequently, accuracy generalizes poorly as genetic distance between training and target sets increases (Beaulieu et al., 2014; Lenz et al., 2017; Resende et al., 2012c). Reduced transferability is typically attributed to a combination of factors, including differences in linkage disequilibrium between markers and causal variants across populations, shifts in allele frequencies and effect sizes, and genuine genotype-by-environment interactions that alter realized genetic architecture across environments (Beaulieu et al., 2014; Lauer et al., 2022; Resende et al., 2012a; Resende et al., 2012b; Slavov et al., 2025). Taken together, these findings motivate prediction frameworks that explicitly model environment-dependent genetic effects rather than assuming that marker effects remain constant across populations and environments.

In our dataset, environmental similarity alone had limited predictive value, but its signal was consistently detectable: among single-predictor models, BIO19 and SPI12_08 performed best, and *PC1*_env_ captured the strongest environmental signal. This indicates that broad-scale hydroclimatic context contributes measurably to crown damage, even if it explains only a small fraction of overall phenotypic variance. Such modest performance of environment main effect prediction models is unsurprising in natural stands (Hennon et al., 2020; Holden et al., 2020), where disease severity integrates fine-scale, spatially structured processes—including local inoculum pressure, microclimatic buffering, stand structure, and host physiological status. These factors are only partially captured by coarse-gridded climate layers (Bütikofer et al., 2020; De Frenne et al., 2021). Nevertheless, even a weak but reproducible environmental signal can be useful in prediction, because it provides a structured proxy for environmental similarity that can help represent environment-dependent departures from genetic main effects.

This weak but reproducible environmental signal became more informative when modelled within a *G×E* framework: explicitly modelling *G×E* in a multi-kernel model yielded small but systematic improvements over genetic main-effect models. The practical importance of this result lies less in increasing peak within-sample accuracy than in reducing the loss of predictive performance when models are transferred across environmental or geographic contexts. In the discovery population, the *G×E* model outperformed the genetic main-effect model across SNP panel sizes, with the largest gains at low marker density (Δ*r* ≈ 0.02–0.03 at 100–200 SNPs) and a smaller but still detectable advantage at 500 SNPs (Δ*r* ≈ 0.015), while differences became minimal at ≥1,000 SNPs. Importantly, the same pattern was observed in the external dataset, where including *G×E* again improved transferability, indicating that the interaction term captured a signal that generalized beyond the discovery population rather than simply increasing within-sample fit. This behaviour is consistent with forest-tree studies showing that multi-environment prediction models that explicitly parameterize *G×E* can retain higher accuracy under across-site or “untested environment” prediction than single-site models, with benefits often strongest for environmentally sensitive growth traits and weaker for wood-quality traits (Beaulieu et al., 2022; Grattapaglia et al., 2018; Li et al., 2017) Variance partitioning provides a mechanistic explanation for why *G×E* improves robustness even when environmental main effects are small. In the discovery set, additive genetic variance dominated (VG = 79.30%), the environmental main effect was negligible (VE = 0.58%), yet the interaction component was clearly non-zero (V*_G×E_* = 14.90%). In the external dataset, the proportional contribution of *G×E* was even larger (V*_G×E_* = 20.87%), while VE remained near zero (0.23%) and additive variance still dominated (V_G_ = 74.78%). This structure—weak shared environment effects but appreciable interaction variance—suggests that broad-scale environmental factors modulate disease outcomes primarily by altering relative genetic performance (genotype-specific sensitivity and/or re-ranking) rather than by imposing a uniform shift across individuals. Similar variance decompositions have been reported in forest trees, where additive effects typically explain most of the variation, but *G×E* contributes a smaller yet consequential component under heterogeneous site conditions (Chen et al., 2019; Dong et al., 2024; Li et al., 2017; Muñoz et al., 2014). In host–pathogen systems, such a pattern is biologically plausible because climatic conditions can influence both pathogen pressure and host stress, while genotypes differ in how strongly those pressures are translated into visible symptoms and crown damage.

The value of modelling *G×E* was most evident under extrapolative validation. For the 500-SNP GWAS-informed panel, predictive ability declined from random CV to LOECO and LOGCO, indicating sensitivity to shifts in environmental and spatial context (internal: *r* ≈ 0.89 in CV vs ≈0.86 in LOECO and ≈0.85 in LOGCO; external: *r* ≈ 0.80 vs ≈0.77 and ≈0.76, respectively). Importantly, incorporating *G×E* mitigated this decline across scenarios, with the largest gain under environmental cluster hold-out (Δ*r* ≈ +0.03; internal LOECO ≈0.89 vs 0.86 for genetic main-effect model), and the same ordering (CV > LOECO > LOGCO) was observed in the external population. This behaviour closely matches patterns reported in forest-tree genomic prediction under “untested environment/site” schemes, where accuracy typically drops most strongly for growth traits but is more stable for wood-quality traits. In white spruce, prediction into untested environments remained moderately high for wood traits (reported correlations ≥0.61) but dropped markedly for growth traits (≥0.24) (Beaulieu et al., 2014). In Norway spruce, cross-site accuracy for tree height declined from 0.76 within-site to 0.44 across sites, whereas the reduction was much smaller for modulus of elasticity (0.64 to 0.59), indicating substantially greater environmental sensitivity of growth-related predictions (Chen et al., 2018). Together, these results suggest that the main value of *G×E* modelling lies less in boosting within-sample fit than in improving robustness when prediction targets fall outside the environmental or geographic envelope represented in the training data—precisely the situation most relevant for deployment across heterogeneous landscapes.

### 4.5 Implications for breeding and conservation

From an applied perspective, our results support genomic prediction as a promising tool for ash breeding, restoration, and conservation, particularly when based on GWAS-informed marker panels and interpreted in an environment-aware framework. The strong additive genetic signal and the high performance of prioritized SNP panels suggest that resilient material can be identified more efficiently than by phenotypic screening alone. At the same time, the improved performance of *G×E*-aware models under extrapolative validation indicates that deployment decisions should account for environmental context—incorporating tools such as climate niche mapping and assisted migration frameworks—rather than relying solely on genomic ranking derived from a single set of sites (Carroll & Boa, 2024; Seidel et al., 2025).

These findings also suggest that moderate, trait-enriched SNP panels may be more useful operationally than very large marker sets, as they captured the bulk of the transferable predictive signal while mitigating the performance decay observed at higher marker densities. More broadly, the polygenic and partly environment-dependent nature of ash dieback resistance argues against management strategies focused on only a few top-ranked genotypes. A more resilient approach is likely to involve the conservation and deployment of genetically diverse material carrying broad standing variation in disease response, thereby supporting both immediate resilience and long-term adaptive potential.

## Acknowledgements

We thank the Forest Gene Bank in Kostrzyca and the local foresters involved in the nationwide survey for identifying candidate ash trees and facilitating field sampling. We also thank Ewa Sztupecka and Katarzyna Meyza for their valuable assistance with sample collection and laboratory work.

## Author Contributions

J.M. and J.B. conceived and designed the study. J.M. and B.U. conducted the fieldwork. B.U. processed raw sequencing data and performed SNP calling. J.M. analyzed the data and wrote the first draft of the manuscript. J.B. acquired funding and supervised the research. All authors contributed to data interpretation, revised the manuscript, and approved the final version.

## Data availability

Raw sequencing reads have been deposited in the NCBI Sequence Read Archive (SRA) and will be made publicly available upon publication. Accession numbers will be provided upon acceptance.

## Supporting information

### Figures

**Figure S1.** Density distribution of the synthetic tree damage index (Syn) among sampled trees. The dashed vertical line indicates the mean Syn value across all individuals.

**Figure S2.** Environmental principal component analysis (PCA) of climatic and drought-related variables.

**Figure S3.** Principal component analysis (PCA) based on genome-wide SNP data. Each point represents an individual.

**Figure S4.** Heatmap of the genomic relationship matrix (GRM) among 320 F. excelsior individuals based on genome-wide SNP data. Colours indicate pairwise kinship coefficients estimated in GEMMA. Hierarchical clustering illustrates relative genetic similarity among individuals.

**Figure S5.** Quantile–quantile (QQ) plot of GWAS p-values for ash dieback severity (Syn) based on the full dataset. Observed versus expected –log10(p-values) under the null hypothesis of no association. The shaded area represents the 95% confidence envelope for uniformly distributed p-values, and the red line indicates the expected distribution.

**Figure S6.** Quantile–quantile (QQ) plot of GWAS p-values for the extreme-phenotype analysis contrasting undamaged (Class 0) and strongly damaged (Class 3) trees. Observed versus expected –log10(p-values) under the null hypothesis of no association. The shaded area represents the 95% confidence envelope for uniformly distributed p-values, and the red line indicates the expected distribution.

### Tables

**Table S1.** Details of sampled *F. excelsior* individuals.

**Table S2.** Pearson correlation coefficients between the disease severity index (Syn) and environmental variables.

**Table S3**. Summary of whole-genome sequencing and alignment statistics.

**Table S4.** SNPs associated with ash dieback severity in the full-dataset GWAS

**Table S5.** SNPs associated with ash dieback severity in the extreme-phenotype GWAS (Class 0 vs. Class 3)

**Table S6.** SNPs with the highest loadings on the first principal component (PC1) in the multilocus PCA of suggestive GWAS loci

**Table S7.** Allele-frequency differences at LD-pruned suggestive GWAS loci between undamaged (Class 0) and strongly damaged (Class 3) ash trees.

**Table S8.** Genotype–phenotype associations for nonsynonymous candidate SNPs at suggestive GWAS loci.

**Table S9.** Summary of the independent external validation set.

