## Supplemental Data 1 for "Environment-aware genomic prediction enhances the transferability of polygenic resistance to ash dieback in *Fraxinus excelsior*"

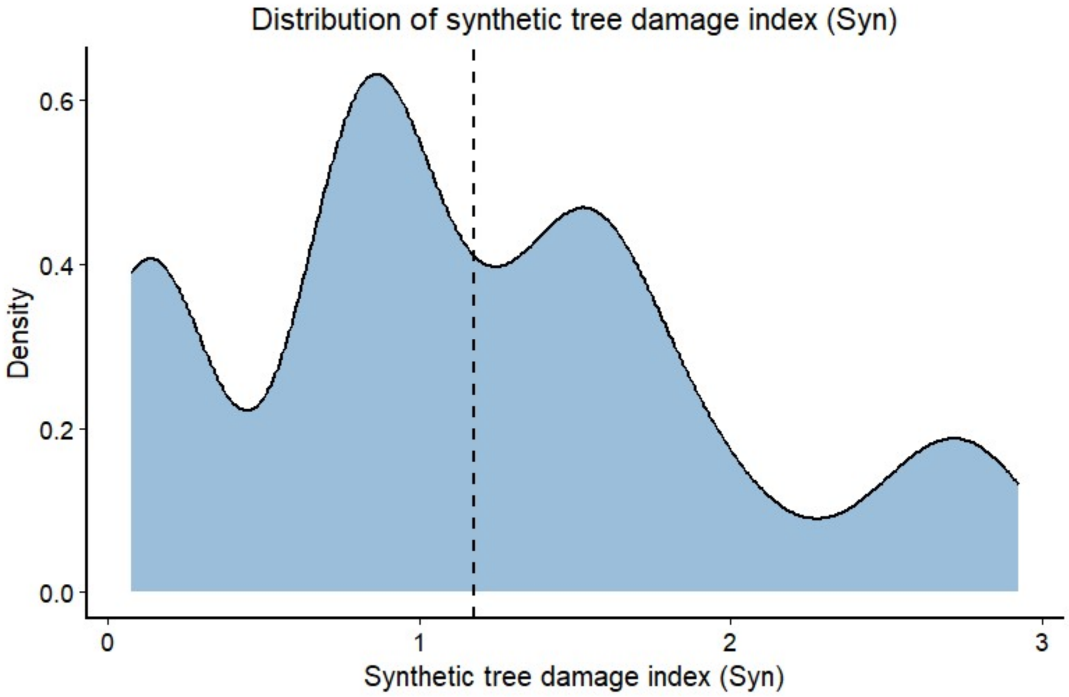


**Figure S1.** Density distribution of the synthetic tree damage index (Syn) among sampled trees. The dashed vertical line indicates the mean Syn value across all individuals.

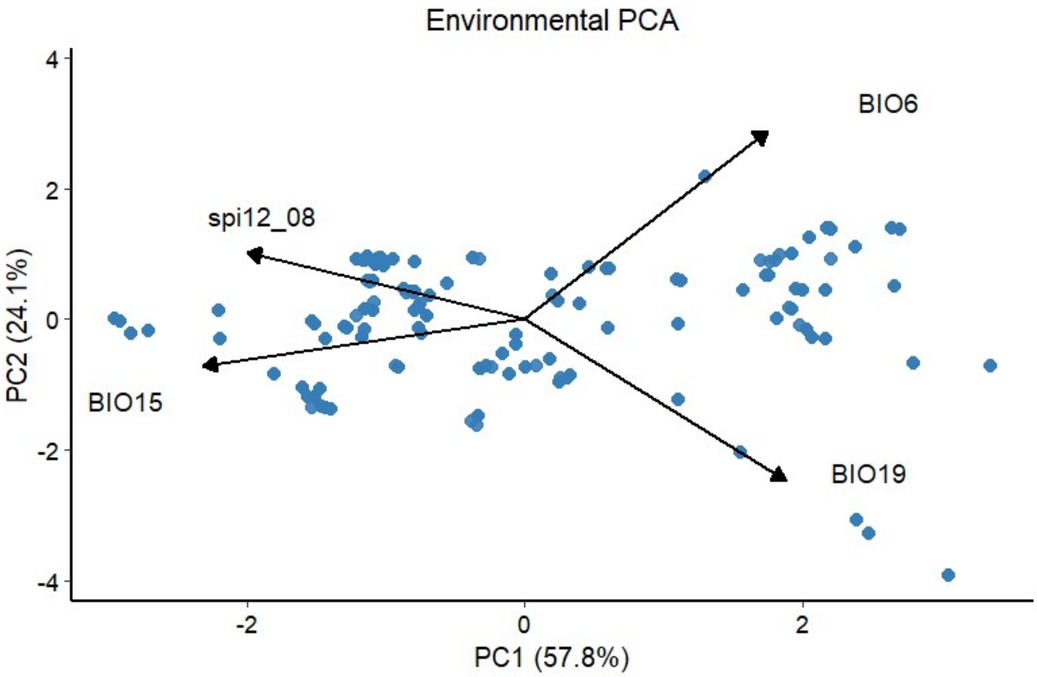


**Figure S2.** Environmental principal component analysis (PCA) of climatic and drought-related variables.

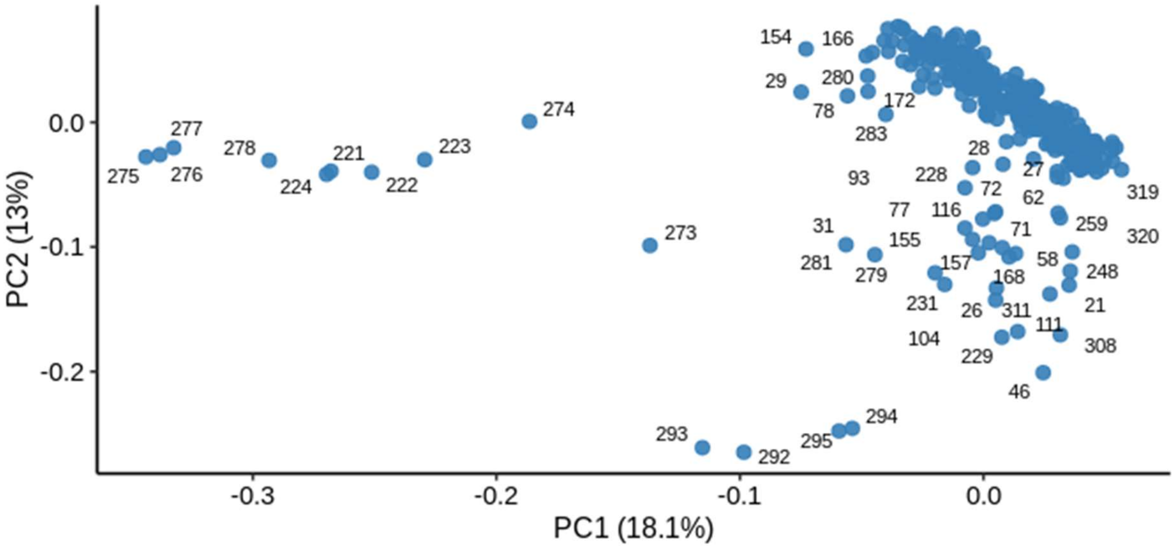


**Figure S3. Principal component analysis (PCA) based on genome-wide SNP data.** Each point represents an individual.

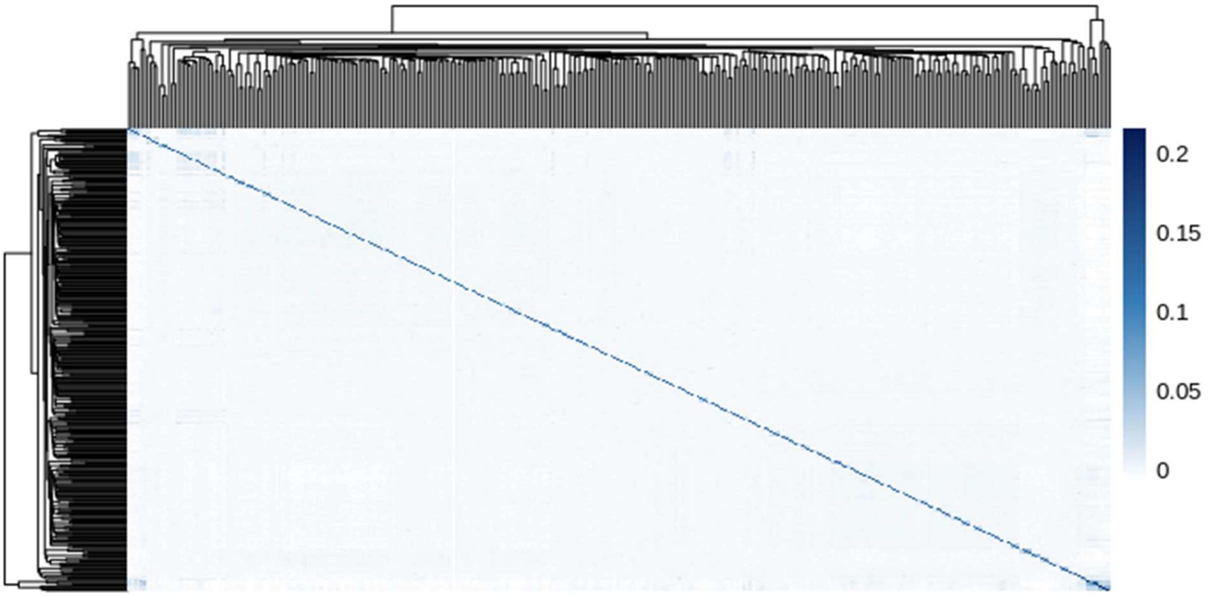


**Figure S4.** Heatmap of the genomic relationship matrix (GRM) among 320 *F. excelsior* individuals based on genome-wide SNP data. Colours indicate pairwise kinship coefficients estimated in GEMMA. Hierarchical clustering illustrates relative genetic similarity among individuals.

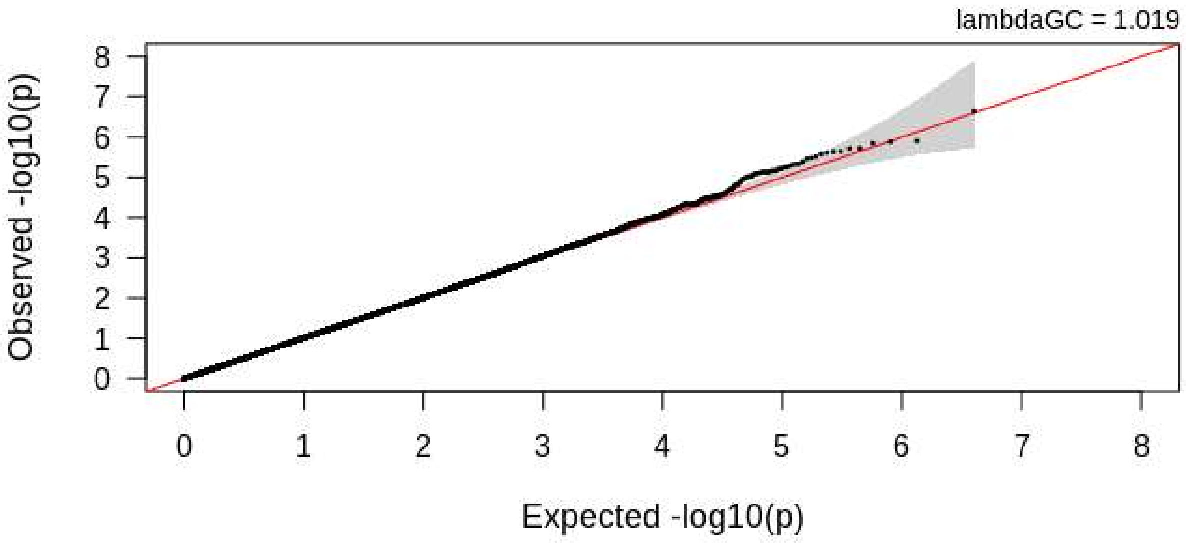


**Figure S5.** Quantile–quantile (QQ) plot of GWAS p-values for ash dieback severity (Syn) based on the full dataset. Observed versus expected –log10(p-values) under the null hypothesis of no association. The shaded area represents the 95% confidence envelope for uniformly distributed p-values, and the red line indicates the expected distribution.

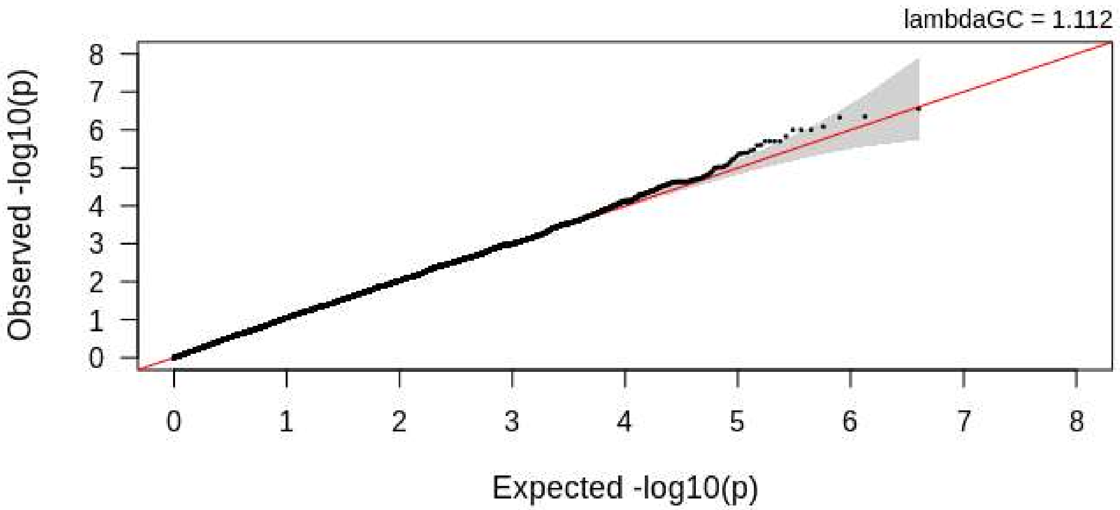


**Figure S6.** Quantile–quantile (QQ) plot of GWAS p-values for the extreme-phenotype analysis contrasting undamaged (Class 0) and strongly damaged (Class 3) trees. Observed versus expected –log10(p-values) under the null hypothesis of no association. The shaded area represents the 95% confidence envelope for uniformly distributed p-values, and the red line indicates the expected distribution.
